# Elimination of senescent cells with senolytic host-directed therapy reduces tuberculosis progression in mice

**DOI:** 10.1101/2025.03.28.645957

**Authors:** Somnath Shee, Yazmin B. Martinez-Martinez, Benjamin Koleske, Shivraj Yabaji, Lester Kobzik, Igor Kramnik, William Bishai

**Author notes:** For correspondence (William Bishai).

## Abstract

By eliciting lung necrosis, which enhances aerosol transmission, *Mycobacterium tuberculosis* (*Mtb*) sustains its long-term survival as a human pathogen. In studying the human-like necrotic granuloma lesions characteristic of *Mtb*-infected *B6.Sst1S* mice, we found that lung myeloid cells display elevated senescence markers: cell cycle arrest proteins p21 and p16, the DNA damage marker γH2A.X, senescence-associated β-galactosidase activity, and senescence-associated secretory phenotype (SASP). These markers were also elevated in *Mtb*-infected aged wild type (WT) mice but not in young WT mice. Global transcriptomics data revealed upregulation of pro-survival (PI3K, MAPK) and anti-apoptotic pathways in *Mtb*-infected *B6.Sst1S* macrophages. As senescent cells are terminally growth-arrested yet metabolically active cells that release tissue-damaging, immunosuppressive SASP, we treated *Mtb*-infected mice with a cocktail of three senolytic drugs (dasatinib, quercetin, and fisetin) designed to kill senescent cells. Senolytic drug treatment prolonged survival and reduced *Mtb* lung counts in *B6.Sst1S* and aged WT mice to a greater degree than young WT mice and concomitantly reduced lung senescence markers. These findings indicate that (1) *Mtb* infection may induce lung myeloid cells to enter a senescent state and that these cells may promote disease progression, and (2) senolytic drugs merit consideration for human clinical trials against tuberculosis (TB).

**Graphical abstract:** 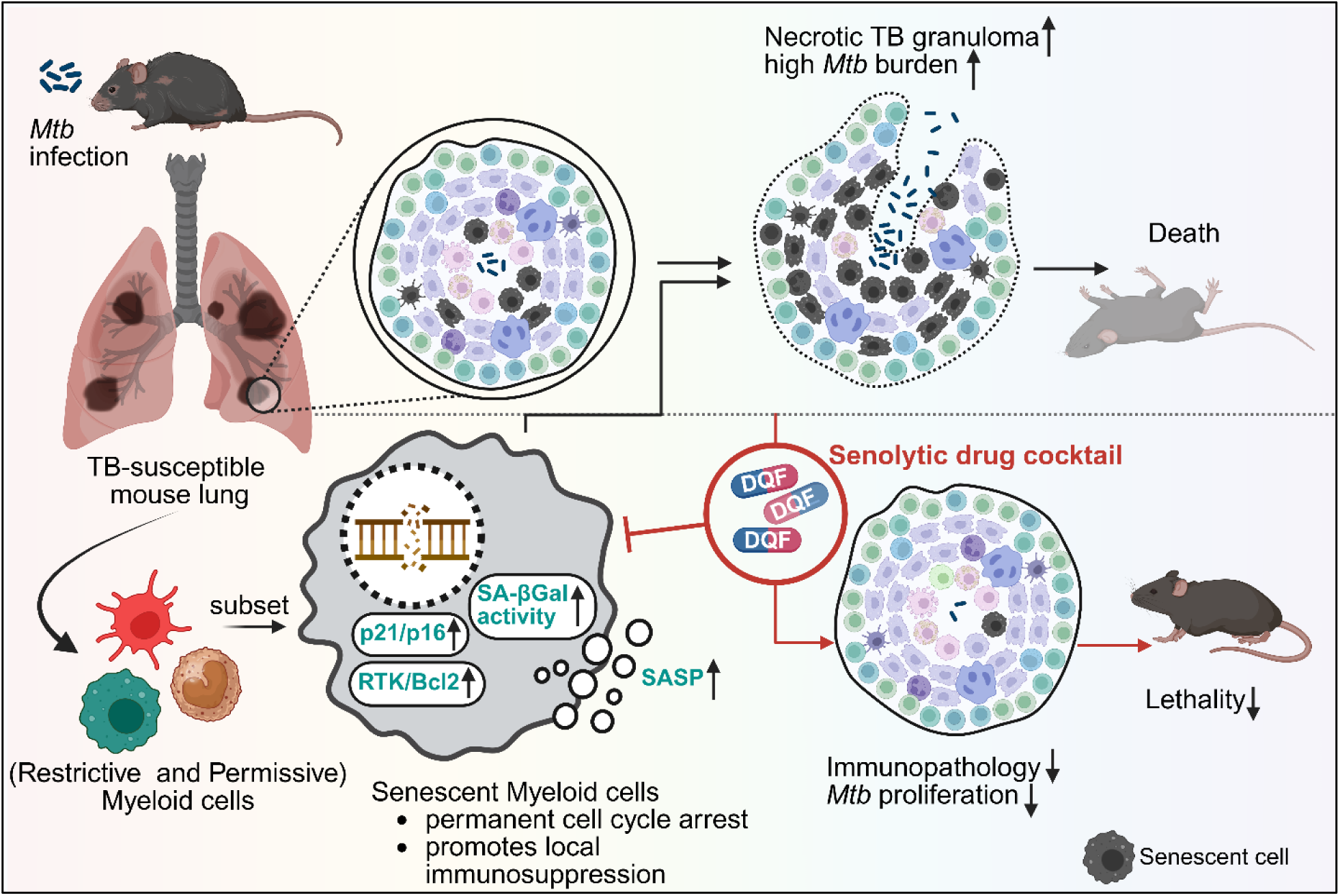

**Highlights:**

- *Mtb* lung infection results in recruitment of both restrictive and permissive myeloid cells to the nascent granuloma.
- *Mtb* infection induces certain permissive myeloid cells to enter a senescent state, characterized by cell cycle arrest and they promote local immunosuppression.
- Treatment with a Senolytic drug cocktail, which kills senescent cells, augments host resistance against *Mtb* proliferation, lethality and immunopathology.

## Introduction

Despite the introduction of 3 new anti-tuberculosis (TB) drugs since 2012, the global TB pandemic remains poorly controlled.^1^ Most humans are resistant to TB and are capable of limiting the bacterial growth and inflammatory damage caused by *Mycobacterium tuberculosis* (*Mtb*) infection.^2–4^ However, susceptible individuals develop active disease characterized by necrotic granulomatous lung lesions that may further transform into lung cavities, leading to lung damage and morbidity even with appropriate anti-TB treatment, and ultimately TB transmission.^5,6^ Understanding the molecular basis for failed granuloma control and necrotic lesion formation is a major unmet need in TB research.

While it is well known that aging is a significant risk factor for progression from contained *Mtb* infection to active TB disease in both high and low-level income countries,^7–11^ there have been relatively few studies investigating the mechanistic basis for age-associated active TB.^12–20^ Senescence is a rapidly expanding research discipline as many investigators have sought to understand the mechanism of gradual tissue failure in age-associated chronic diseases such as dementia, heart failure, arthritis, and renal failure.^21^ The accumulation of senescent cells is a principal mechanism of functional tissue impairment in aging.^22,23^ A key lesson from the senescence literature is that aged tissues are populated by long-lived, non-dividing, functionally impaired senescent cells. These cells release deleterious signaling molecules-collectively termed senescence-associated secretory phenotype (SASP), that locally contribute to immunosuppression, dysregulated inflammation, and tissue failure.^24–27^ Indeed, transplantation of isolated senescent cells led to premature aging in mice.^28,29^ In this context, small molecule senolytic drugs—designed to induce apoptosis in the non-dividing senescent cells—prolong murine lifespan, and improve organ function in humans with chronic disease.^30,31^ To gain insight into the factors that lead to the loss of immune containment of *Mtb*, we study the *B6.Sst1S* mice, which develop human-like necrotic lung granulomas following *Mtb* infection in contrast to most wild type (WT) mouse strains.^32–38^ These mice harbor the *Sst1S* allele, which has loss of function mutations in the Sp140 chromatin remodeling protein that plays a key role in immune cell gene regulation.^39,40^ Additionally, the closely related *Sp140^-/-^* mouse strain was characterized to exhibit a population of interstitial macrophages (IM) and plasmacytoid dendritic cells (pDC) that aberrantly secrete type I IFN (associated with TB exacerbation),^41–43^ are poorly responsive to IFNγ, and may play a key role in granuloma breakdown in these mice.^44^

In this paper, we report that a distinct population of lung myeloid cells in *Mtb*-infected *B6.Sst1S* mice display higher levels of senescence markers prior to necrotic granuloma formation and are subsequently found in close association with non-controlling, early necrotic granulomas. Similar senescence marker increases are present in aged, *Mtb*-infected WT C57BL/6 (B6) mice, but not young WT B6 mice. Treatment with a three-drug cocktail of FDA-approved senolytic drugs to eliminate these senescent cells restored containment of *Mtb* proliferation, prolonged mouse survival, and reduced immunopathology and senescence markers. Thus, during TB infection a population of lung myeloid cells enters a senescent state and may contribute to failure of immune containment and progression to necrosis.

## Results

### Elevated senescence markers were detected in BMDMs from *B6.Sst1S* and aged WT B6 but not young WT B6 mice *in vitro*

We previously observed that TNFα exposure induced a stress response characterized by overexpression of IFNβ (Figure S1A) and the integrated stress response (ISR) in *B6.Sst1S* bone marrow-derived macrophages (BMDMs).^37,38^ Reanalysis of transcriptomic signatures in uninfected *B6.Sst1S* versus WT B6 BMDMs with and without TNFα treatment (GSE99456)^38^ suggested upregulation of cellular senescence pathways, enhanced DNA damage responses, activation of mitotic checkpoints, pro-survival ERK1/ERK2 protein kinase cascade, and cellular response to IFNβ (by Metascape Analysis;^45^ Figure S1B-S1D, Supplementary File S1) in *B6.Sst1S* BMDMs. Because chronic ISR activation and heightened Type I interferon exposure can be a potential trigger of cellular senescence,^46–50^ we tested uninfected TNFα-stimulated BMDMs from young *B6.Sst1S* mice (8-12 weeks old) alongside BMDMs from young and aged (>72 weeks old) WT B6 mice for the presence of canonical senescence markers (Figure 1A).^51–54^ *B6.Sst1S* and aged WT B6 BMDMs showed greater activity of senescence-associated β-galactosidase (SA-βGal) than young WT B6 BMDMs, although not to the level seen with etoposide, an inducer of DNA damage-mediated senescence (Figure 1B-1D, Figure S2A). In addition, we observed the induction of SASP genes (*Il1β, Il1α, Il6, Mmp12, Mmp13, Ccl2, Ccl8, Pai1*) transcription in TNFα-treated *B6.Sst1S* BMDMs compared to similarly treated WT B6 cells (Figure 1E). Moreover, TNFα-treated *B6.Sst1S* BMDMs and aged WT B6 BMDMs showed the presence of established biomarkers of senescence: higher levels of the cyclin-dependent kinase inhibitor P16 (cell cycle arrest), higher phospho(S139)-H2A.X (DNA damage marker), lower Lamin B1 (structural constituent of the nuclear lamina, Figure S2B), and reduced cell proliferation as measured by BrdU-incorporation assay (Figure 1F, S2C) without a significant reduction in viability (Figure S2D). BMDMs from young *B6.Sst1S* and aged WT B6 mice (the same cells with higher senescence markers following TNFα induction) showed poorer *Mtb* containment than those from young WT B6 (Figure 1G). In sum, in line with the transcriptomics data, TNFα-treated *B6.Sst1S* BMDMs indeed exhibited numerous markers of senescence similar to BMDMs from aged WT B6 mice or etoposide-treated WT B6 controls.

**Figure 1.**
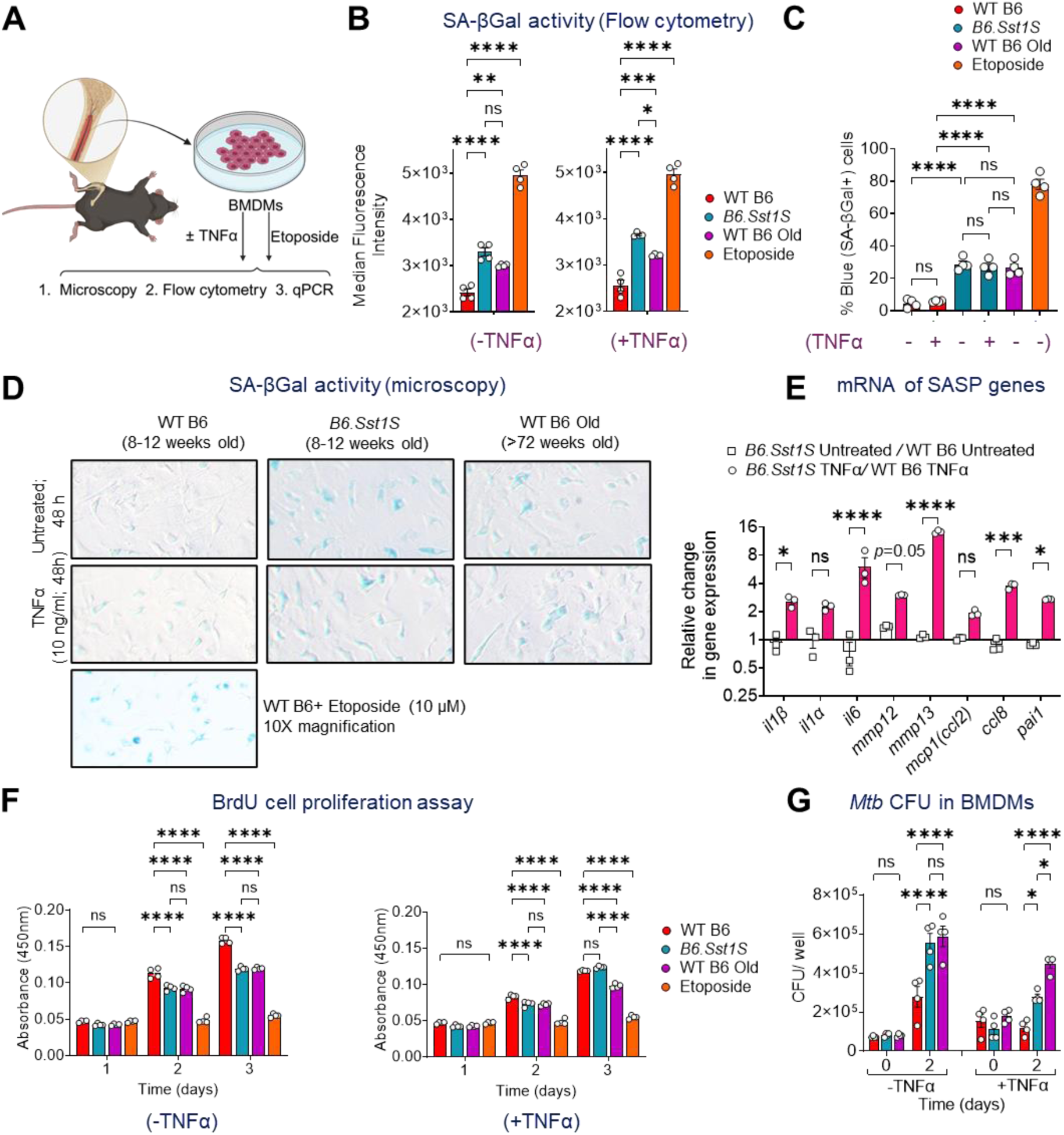
Evidence of senescence-like phenotype in uninfected BMDMs derived from young *B6. Sst1S* and WT B6 Old mice. (A) Schematic to determine senescence associated markers in BMDMs. (B) Median Fluorescence intensity of fluorescent SA-βGal substrate by live BMDMs in presence or absence of TNFα for 48 h. 10 µM Etoposide treated WT B6 BMDMs or BMDMs from aged mice are used as positive control. (C,D) quantification and representative images of SA-βGal-positive BMDMs (PFA-fixed) located at the center of the TC-treated well at 20X magnification by bright field microscopy. (E) mRNA levels of SASP-associated genes normalized to ActB as determined by qPCR. (F) BrdU incorporation by BMDMs in absence or presence of TNFα, respectively with time. (G) BMDMs were infected for 4 hours with *Mtb* H37Rv at MOI=10, followed by removal of extracellular bacteria by extensive washing with warm media. Intramacrophage survival was determined by enumerating CFU at 2 days post-infection. The data are means ± SEM. Data is representative of two independent experiments done with n≥2. (*p*>0.05: ns, *p*<0.05: *, *p*<0.01: **, *p*<0.001: ***, *p*<0.0001: ****, one-way ANOVA with Tukey’s (B, C, F, G) or two-way ANOVA with Sidak’s (E) multiple comparisons test).

### Senescence markers are elevated in non-controlling lung lesions of *B6.Sst1S* mice and in lung myeloid cells from *Mtb*-infected young *B6.Sst1S* and aged WT B6 mice

To examine whether senescence signatures appear in the context of active lung infection, we performed spatial transcriptomics to profile neighboring cells associated with controlling (C) versus non-controlling (NC) lesions in *B6.Sst1S* mouse lungs (Figure 2A). To do this, we first administered 1×10^6^ CFU of *Mtb* subcutaneously in the hock of *B6.Sst1S* mice and allowed generation of chronic lesions in a post-primary pulmonary TB model.^55^ Lung samples were harvested at 14 weeks post-infection and subjected to unbiased spatial transcriptomics targeting cells near lesions. We categorized the lesions into non-controlling (NC; advanced multibacillary lesions with necrotic cores) and controlling granulomas (C; non-necrotic paucibacillary lesions), as determined by acid-fast and H&E staining (Figure 2A). We identified 734 differentially expressed genes (DEGs; log_2_fold change>1 or <-1, adjusted *p*< 0.05) in the neighboring cells from NC vs C lesions (Figure 2B, Supplementary File S2). Functional pathway enrichment analysis^45^ identified inflammatory response, extracellular matrix organization, lung fibrosis and negative regulation of cell proliferation pathways to be significantly higher in NC lesions (Figure S3A). Moreover, there was a statistically significant overlap between NC vs C lesion-DEGs and the SenMayo^56^ senescence-associated gene set (Fisher’s exact test, two-sided, *p*< 0.0001; Figure S3B, S3C) indicating enrichment of senescence-associated genes in non-controlling lung lesions. These spatial transcriptomics data indicate that lung cells expressing senescence markers are uniquely present near and within granulomas progressing to necrosis in *Mtb*-infected *B6.Sst1S* mouse lungs.

**Figure 2.**
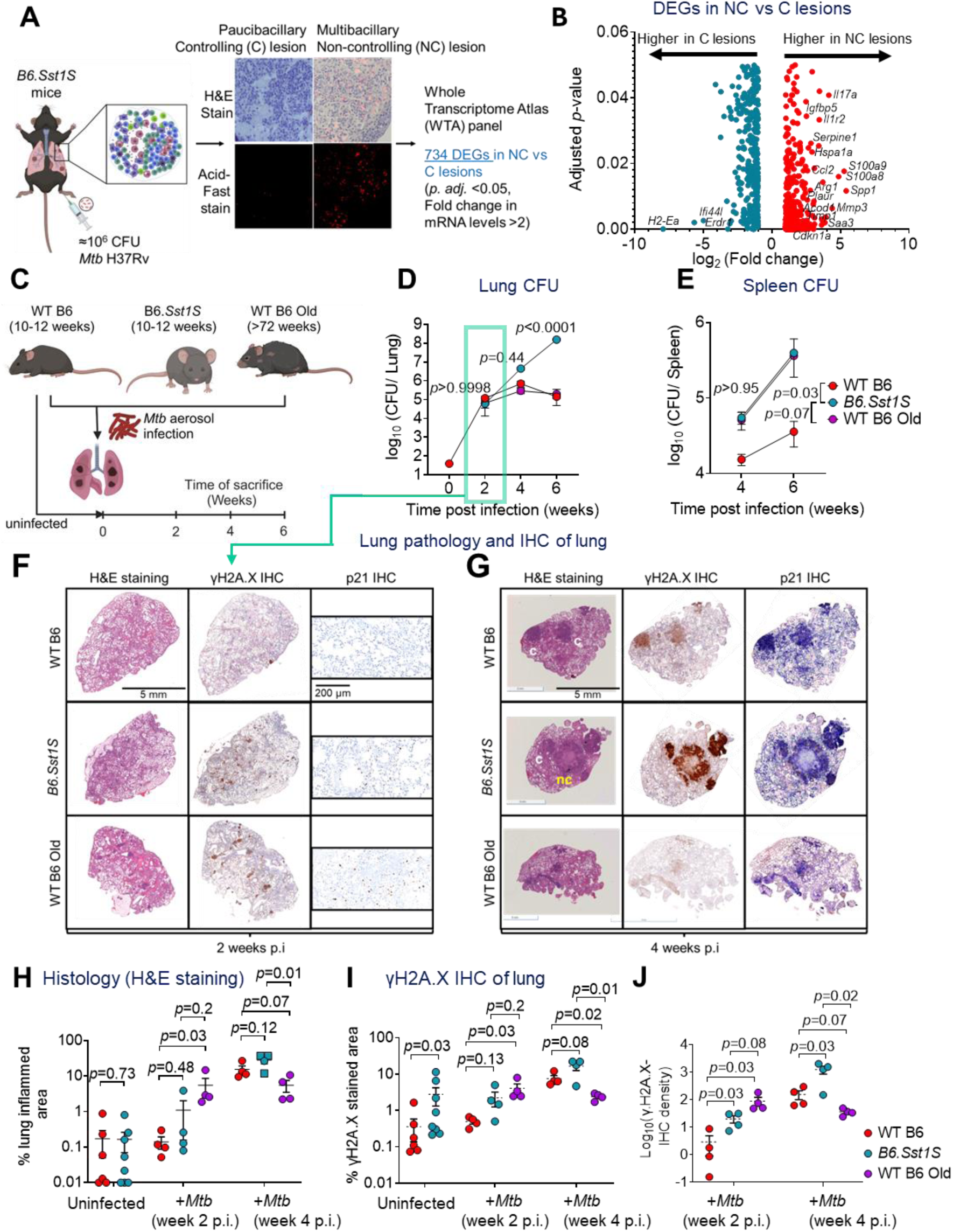
Detection of Senescence-associated markers during early stages of disease progression in *Mtb* infected mice. (A) Schematics of Spatial transcriptomics using a Nanostring GeoMX Digital Spatial Profiler to determine differential gene expression in cells at non-controlling multibacillary granulomatous lesions (*NC*) relative to controlling lesions (*C*) in *B6.Sst1S* mice lungs. (n= 8 ROI). (B) Volcano plot to show differentially expressed genes between controlling and non-controlling granulomatous lesions. Significantly upregulated (fold change> 2) and downregulated (fold change< 2) genes (*p.adj*< 0.05) in *NC* relative to *C* granulomas are shown in red and cyan, respectively. Top senescence associated genes in upregulated DEGs are labelled. (C) Schematics of the experimental strategy to measure senescence markers *in vivo* at 2- and 4-weeks post infection. Day1 CFU=39. *Mtb* H37Rv burden at indicated time points in WT B6, *B6.Sst1S* and WT B6 old mice (D) lungs and (E) spleen. The data are means ± SEM. *p* was determined by two-way ANOVA with Tukey’s multiple comparisons test (*p*>0.05: ns, *p*<0.05: *, *p*<0.0001: ****). Representative H&E-stained and Immunohistochemistry against γH2A.X and p21 of mice lungs at (F) 2 weeks p.i. (arrow indicates no difference in *Mtb* burden) and (G) 4 weeks p.i. (H) ImageJ quantification of lung pathology (H&E-stained area) in uninfected mice and *Mtb* H37Rv infected mice lungs at 2 weeks and 4 weeks p.i. (n=4-6). Squares indicate necrotic granulomas. (I) ImageJ quantification of γH2A.X-stained area in uninfected mice and *Mtb* H37Rv infected mice lungs, and (J) corresponding γH2A.X-IHC intensity at 2 weeks and 4 weeks p.i. (n=4-6 mice/group). The data are means ± SEM. Each data point represents a mouse. Statistical analysis between two groups was measured by unpaired two-tailed Student’s t test (normal distribution) or Mann Whitney test.

To examine the accumulation of senescence markers *in vivo*, we infected young *B6.Sst1S*, young WT B6 mice, and aged WT B6 mice with *Mtb* by the aerosol route (Figure 2C). *Mtb* proliferation was significantly higher uniquely in young *B6.Sst1S* mice, with necrosis appearing as early as 4 weeks post-infection (wpi) (Figure 2D, 2G). Both young *B6.Sst1S* and aged WT B6 showed increased dissemination to the spleen at 6 wpi, suggesting that aged WT B6 mice have a defect in containment compared to young WT B6 (Figure 2E). At 2 wpi, when lung CFU counts were equivalent across all three groups, we observed a higher degree of immunopathology as measured by area of lung consolidation in the H&E-stained lungs of aged WT B6 compared to young WT B6 (*p*= 0.03), while young *B6.Sst1S* mice showed similar degree of lung disease (*p*=0.48, Figure 2F, 2H). At 4 wpi the degree of pathology in aged WT B6 mice lungs remained the same as at 2 wpi, while there was considerable progression in young *B6.Sst1S* and young WT B6 (Figure 2G, 2H).

Next, to extend our *in vitro* observations with BMDMs, we aimed to determine if markers of senescence were similarly elevated during *in vivo* mouse infections. We evaluated lung levels of γH2A.X and cyclin-dependent kinase inhibitor p21 by immunohistochemistry at the 2 wpi timepoint, when no differences in CFU and histopathology were present (Figure 2D, 2F, 2H). Interestingly, we observed that both young *B6.Sst1S* and aged WT B6 mice showed qualitatively higher immunostaining for γH2A.X and p21 by IHC (Figure 2F), and these levels continued to increase at 4 wpi in *B6.Sst1S* mice but not aged WT B6 (Figure 2G). Quantification of the γH2A.X-stained area and γH2A.X staining-intensity of the lung confirmed significantly elevated levels for both *B6.Sst1S* and aged WT B6 mice relative to young WT B6 mice at 2 wpi (Figure 2I, 2J). To refine this analysis, we turned to flow cytometric measurements of widely accepted senescence markers (p21, p16 and SA-βGal)^53,57^ with lung and spleen cells (Figure 3A, Figure S4). To examine the temporal relationship (cause or effect) between senescence and elevated *Mtb* burden, we measured the senescent cell populations at an early time point, 10 days post-infection (dpi) when lung CFU burdens are the same across all groups (Figure 3B, 3C, Figure S5A). We found that the percent double positive p16^+^p21^+^ cells and the percent SA-βGal^+^ cells out of all live cells were significantly higher (≈2-3-fold) in the lungs of *Mtb*-infected young *B6.Sst1S* and aged WT B6 mice than in young WT B6 at 10 dpi (Figure 3D, 3E). Senescence markers in spleen cells were not elevated in any group at this 10 dpi (Figure 3D, 3E), likely because there was little or no *Mtb* dissemination to spleen (Figure 3C). Importantly, myeloid cells— predominantly alveolar (AM) and interstitial macrophages (IM) accounted for the majority of the enrichment in senescence markers (double positive p16^+^p21^+^ cells and SA-βGal^+^ cells) as seen in young *B6.Sst1S* and aged WT B6 mice lung cells (Figure 3F, 3G). No differences in senescence marker abundance between the three types of mice were observed for B cells, and with T cells we found a modest elevation of SA-βGal^+^ senescence marker in aged WT B6 mice which was not seen with the double positive p16^+^p21^+^ marker (Figure 3F, 3G). Similar data were observed at 20 dpi (Figure S5B, S5C). Lastly, we evaluated lung levels of SASP cytokines in mice lungs. We did not detect enrichment of cytokine levels in mouse lungs at 10 dpi, except IL-17a in aged WT B6 (Figure S5D). However, five of the eight tested (IL-1β, IL-6, TNFα, IL-17a, and GM-CSF) were significantly elevated in young *B6.Sst1S* mouse lungs compared to young WT B6, although no differences were seen in aged WT B6 at 4 wpi and 20 dpi (Figure 3H). Thus during *in vivo* mouse infections, certain senescence markers—most notably macrophages doubly positive p16^+^p21^+^, SA-βGal^+^, and overall γH2A.X levels--are elevated in young *B6.Sst1S* and aged WT B6 mice compared to young WT B6 mice, at a time point where no differences in CFU or histopathology are detected. These data indicate that emergence of senescent cells precedes the failure of *Mtb* containment and progression to necrosis in the TB-susceptible mice.

**Figure 3.**
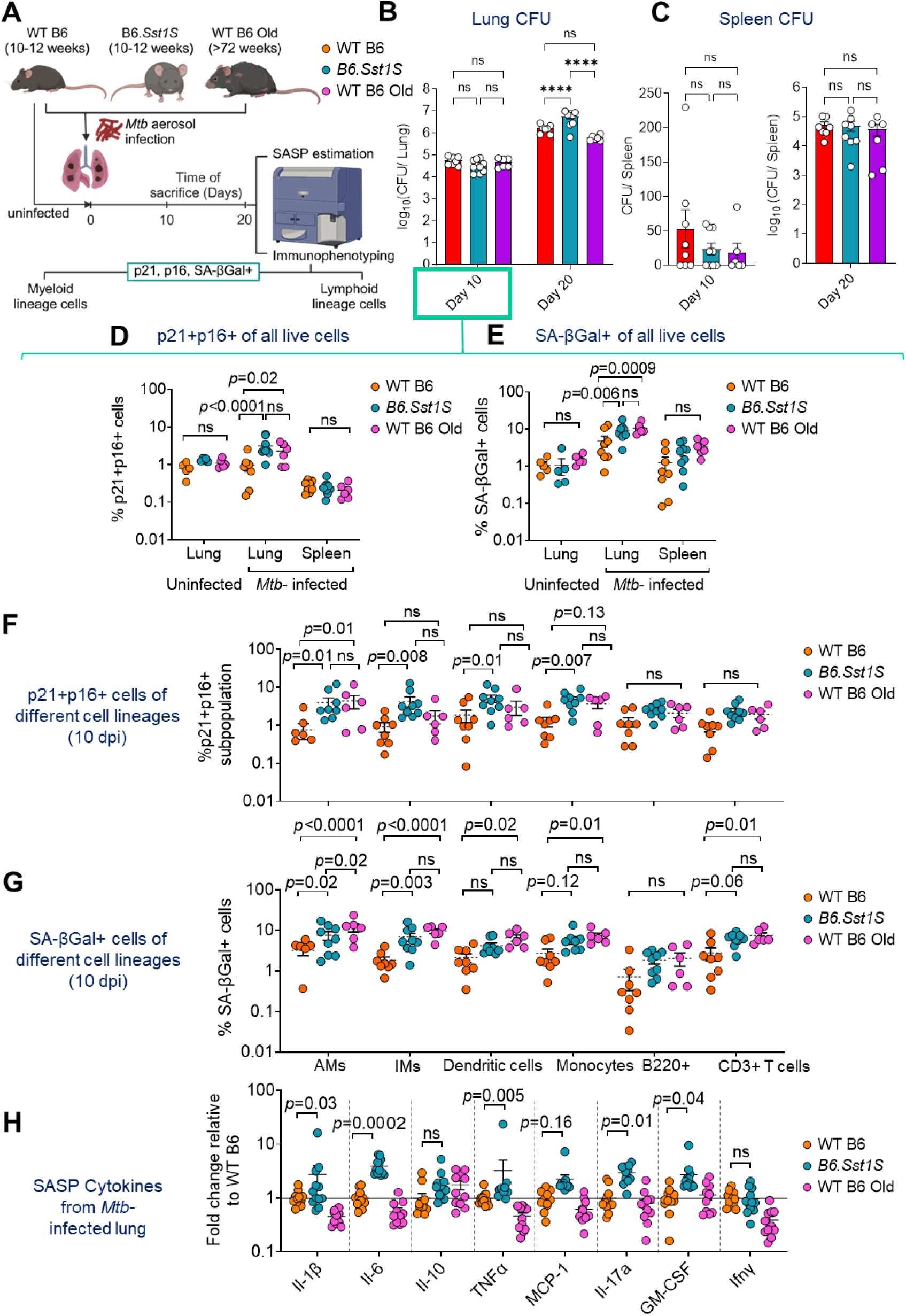
Detection of senescent cell population and SASP cytokines by flow cytometry during early stages of disease progression in *Mtb-*infected mice. (A) Schematics of the experimental strategy to measure senescence markers (p21, p16 and SA-βGal activity) and SASP cytokines levels in mice lungs at 10 and 20 days p.i. Young WT B6, *B6.Sst1S*, and old WT B6 mice were aerosol-infected with 55 CFU of *Mtb* H37Rv. *Mtb* H37Rv burden in mice (B) lungs and (C) spleen are shown at indicated time points (n= 6-9 mice/ group). The data are means ± SEM. Each data point represents a mouse. Statistical analysis was calculated by two-way (B) or one-way (C) ANOVA with Tukey’s multiple comparisons test. (*p*>0.05: ns, *p*<0.0001: ****). (D) %p21+p16+ and (E) %SA-βGal+ (CellEvent Senescence green+) cells out of all live lung (uninfected and *Mtb-* infected mice) and Spleen (*Mtb-*infected mice) cells at 10 days p.i. as determined by multicolor flow cytometry. (F) %p21+p16+ and (G) %SA-βGal+ cells out of different lung-cell types at 10 days p.i., as determined by multicolor flow cytometry. Gating for p16⁺, p21⁺, and SA-βGal⁺ events for each cell type were established using Fluorescence minus one (FMO) controls and reference cells from *Mtb*-infected WT B6 mice (set at 1%). (AMs: Alveolar macrophages, IMs: Interstitial macrophages) **(H)** Normalized concentration of SASP-cytokines in lung homogenates at 20 dpi and 4 wpi relative to WT B6 mice as measured by a LEGENDplex™ Mouse Inflammation Panel. The data are means ± SEM. Each data point represents a mouse (n=11-12). (two-way ANOVA with Tukey’s (D-G) or Dunnett’s (H) multiple comparisons test).

### ‘Omics studies show upregulation of the phosphatidylinositol 3-kinase (PI3K) and the ERK1/2 cascades which are pro-survival, as well as certain anti-apoptotic pathways in *Mtb*-infected BMDMs from *B6.Sst1S* mice

Senescent cells characteristically express cell cycle arrest and pro-inflammatory transcriptional profiles^53,54,58^. To compare the gene expression in stressed *B6.Sst1S* relative to young WT B6 BMDMs, we performed bulk RNA-sequencing with cells subjected to TNFα stress, *Mtb* infection, or both (Figure 4A). BMDMs were exposed to (1) only TNFα (10 ng/ml; 48 h), (2) only *Mtb* infection (MOI=1, 24 h), or (3) 10 ng/ml TNFα pretreatment, followed by *Mtb* infection in the continued presence of TNFα (Figure 4A). Despite increased *Mtb* proliferation in TNFα-treated *B6.Sst1S* BMDMs, cell viability remained intact across all groups (Figure S6A, S6B). When treated with “TNFα only” for 48 h, we identified 1116 DEGs (*p.adj.<* 0.05, log2fold change >1 or <-1) with 597 upregulated and 519 downregulated genes in *B6.Sst1S* relative to similarly treated WT B6 BMDMs (Supplementary File S3). In line with previous data, gene ontology (GO) analysis for biological processes identified-cellular response to interferon-β (GO:0035458; *p.adj*.< 1e-9), defense response to virus (GO:0051607; *p.adj*.< 1e-9), cellular response to interferon-gamma (GO: 0071346; *p.adj*.= 2e-5), and innate immune response (GO: 0045087; *p.adj*.= 1e-5) to be significantly enriched in the *B6.Sst1S* BMDMs (Figure 4B, Supplementary File S3). Interestingly, a distinct transcriptome was observed in *Mtb*-infected *B6.Sst1S* BMDMs (MOI=1; 24 h) (Figure 4B) characterized by 940 DEGs (*p.adj.<* 0.05, log_2_fold change >1 or <-1) with 455 upregulated and 485 downregulated genes in *B6.Sst1S* relative to similarly treated WT B6 BMDMs (Supplementary File S4). Gene ontology (GO) analysis for biological processes identified positive regulation of epithelial cell proliferation (GO:0050679; *p.adj.*= 7.97e-6), positive regulation of PI3K signaling (GO:0014068; *p.adj.*= 1.6e-5), angiogenesis (GO:0001525; *p.adj.*= 1.6e-5), positive regulation of IL-6 production (GO:0032755; *p.adj.*= 0.007), and negative regulation of extrinsic apoptotic signaling (GO:2001240; *p.adj.*= 0.007) to be significantly enriched in the *Mtb*-infected *B6.Sst1S* BMDMs (Figure 4B, Supplementary File S4). The third group (exposure to both Tnfα and *Mtb*) showed 1177 DEGs (*p.adj.<* 0.05, log_2_fold change >1 or <-1) with 647 upregulated and 530 downregulated genes in *B6.Sst1S* relative to similarly treated WT B6 BMDMs (Supplementary File S5). Gene ontology (GO) analysis for biological processes identified receptor tyrosine kinase (RTK) signaling pathway (GO:0007169; *p.adj.*= 0.006), positive regulation of PI3K signaling (GO:0014068; *p.adj.*= 0.001), angiogenesis (GO:0001525; *p.adj.*= 6.34e-5), positive regulation of cell proliferation (GO:0008284; *p.adj.*= 0.001) and apoptotic processes (GO:0043065; *p.adj.*= 0.03), neutrophil chemotaxis (GO:0030593; *p.adj.*= 0.003), and positive regulation of ERK1 and ERK2 cascade (GO:0070374; *p.adj.*= 0.005) to be significantly enriched in the *B6.Sst1S* BMDMs (Figure 4B, 4C, Supplementary File S5). As expected, the genes absent in *B6.Sst1S* cells (*Sp140, Sp110*) were among the top downregulated genes in *B6.Sst1S* BMDMs (Figure 4C). Interestingly, we observed cyclin-dependent kinase inhibitor *Cdkn1c,* senoantigen *Gpnmb*^59^ and several genes included in SenMayo^56^ gene set (*Arg2, Ptger2, Iqgap2, Selplg, Mmp10, Mmp12, Gem, Ets2*), to be upregulated in *B6.Sst1S* BMDMs (Figure 4C). Therefore, stressed *B6.Sst1S* BMDMs exhibited upregulation of pro-survival pathways (PI3K-ERK1/2/MAPK cascade), enriched SASP production (Metascape analysis; Log(*q*-value)= −15.49), and SASP expression regulatory pathways (RTKs), which are characteristic of senescent cells. Additionally, these cells showed increased expression of anti-apoptotic genes (Bcl2a1b, Bcl2a1d, Itga6, Tnfaip8), suggesting resistance to apoptosis. Together, these changes suggest that *Mtb*-infection and/or TNFα stress promotes macrophage entry into a long-lived and non-dividing state characteristic of senescence.

**Figure 4.**
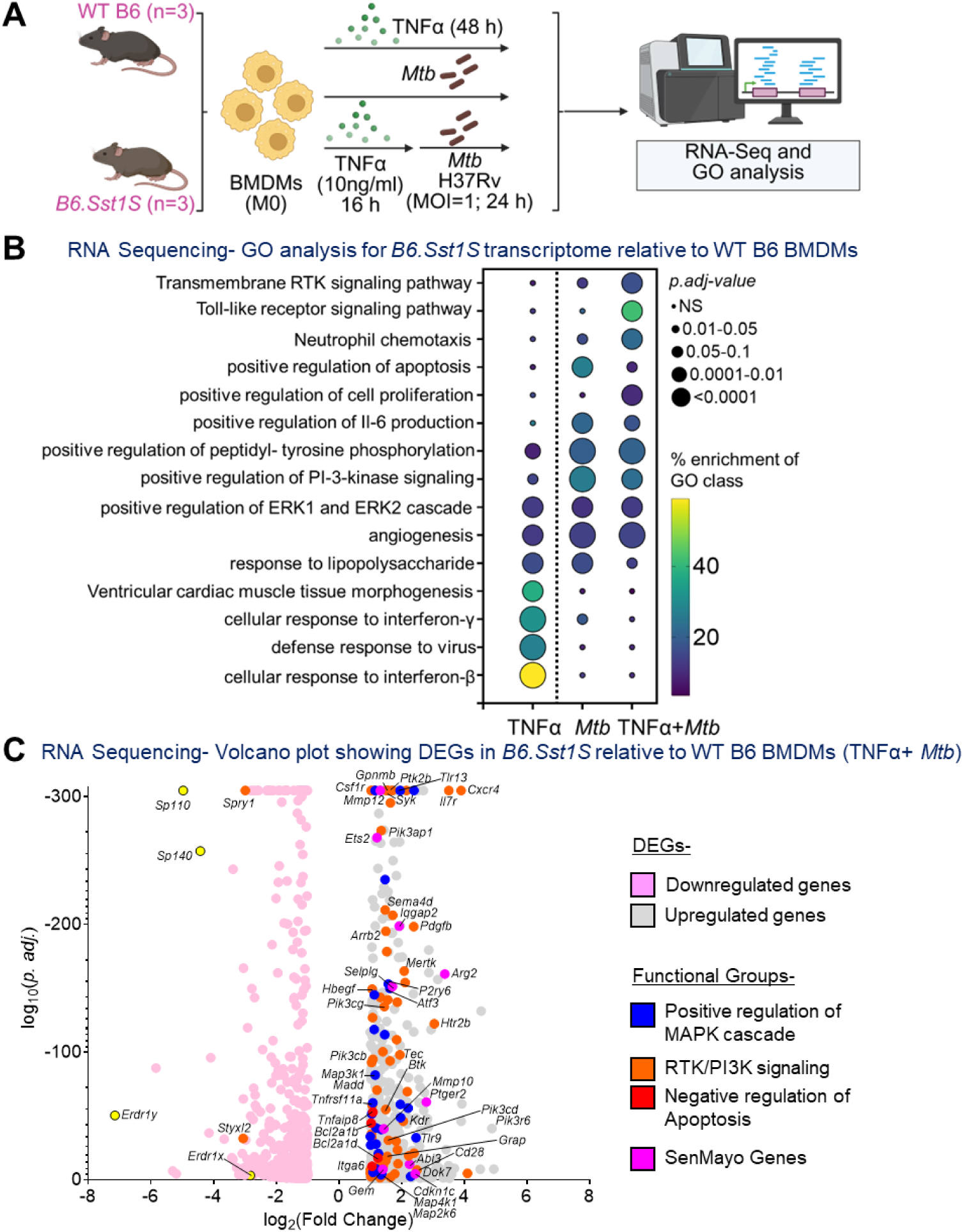
Global transcriptomics data by RNA Sequencing revealed upregulation of pro-survival pathways-PI3K-ERK1/2 and MAPK cascade in BMDMs derived from young *B6.Sst1S.* (A) Schematic of the experiment. (B) Selective representation of Gene-Ontology (GO) analysis for the differentially expressed genes indicating all enriched (*p.adj*< 0.05) signaling pathways under TNFα, *Mtb* and TNFα+ *Mtb-* treatment conditions. The size of the bubble indicates *p.adj* values relative to similarly treated WT B6, and color indicates the percentage of DEGs out of total genes in that GO class. (C) Volcano plot of DEGs (=1177) with upregulated genes in TNFα+ *Mtb-treated B6.Sst1S* BMDMs (log2 fold change >1; *p. adj.<* 0.05) shown in light gray and downregulated genes (log2 fold change <1; *p. adj.*< 0.05) shown in light pink, relative to stressed WT B6 BMDMs. Genes representative of top significantly enriched GO class-Receptor tyrosine kinase signaling/ PI3K pathway (shown in orange), MAPK cascade (blue), anti-apoptotic genes (red) and SenMayo gene set (pink) are shown. Top expected downregulated genes-Sp140, Sp110 and Erdr1 are shown in yellow.

### Therapy with senolytic drugs which kill senescent cells improves host survival during *Mtb*-infection in *B6.Sst1S* mice

A potential strategy to combat aging has been to employ senolytic drugs designed to kill senescent cells and reverse their SASP-mediated local deleterious effects. Indeed, several senolytic drugs have been shown to prolong lifespan in rodent models,^60^ and reduce age-associated disease morbidity in human clinical trials.^21,58,61–63^ We selected a cocktail of three of the best-characterized senolytic agents: dasatinib (D), quercetin (Q), and fisetin (F).^64^ Dasatinib is a tyrosine kinase inhibitor active against Src family and other tyrosine kinases.^65^ Inhibiting multiple kinases promotes apoptotic cell death in senescent cells by disrupting pro-survival signaling networks such as the PI3K, ERK1/2, and other MAPK pathways.^60^ Quercetin and fisetin are flavonoid antioxidants found in common fruits and vegetables; they inhibit both the anti-apoptotic BCL-2 pathway and the pro-survival PI3K and ERK1/2 pathways.^66–69^ Because these pathways are up-regulated in *Mtb*-infected *B6.Sst1S* macrophages (Figure 4B,4C), we reasoned that selectively targeting senescent myeloid cells within TB lung lesions with the DQF cocktail would contain intracellular *Mtb* growth.

First, we confirmed that D, Q, and F do not exert direct anti-microbial activity against *Mtb* at pharmacologically relevant concentrations (Figure S7A, S7B). We then evaluated the host-directed effect of DQF alone or DQF in combination with partial antimicrobial therapy (comprised of ethambutol and rifampicin (DQF + Eth/Rif + Eth)) following an aerosol *Mtb* infection that implanted 91 CFU on day 1 (Figure 5A, 5B). Our rationale for co-administering antimicrobials was to prolong mouse survival giving a longer time window to observe the impact of the host-directed drugs. One week of rifampin (bactericidal) was intended to bring lung CFU counts down to ∼6 log_10_ units, and 4 weeks of ethambutol (bacteriostatic) was intended to extend survival with stable high lung CFU burden. Three weeks post-infection, when the lung CFU burden was 7.18 log_10_CFU units (Figure 5B), young *B6.Sst1S* mice were randomly apportioned to four oral gavage treatment groups for a period of 28 days: (i) vehicle, (ii) DQF (d21-49), (iii) Eth/Rif (d21-28) + Eth (d29-49) and (iv) DQF (d21-49) +Eth/Rif (d21-28) + Eth (d29-49) (Figure 5A). Compared with vehicle, mice receiving DQF alone showed a trend towards prolonged survival in a subset of mice (*p*= 0.09) with a certain degree of weight stabilization (Figure 5C, 5D). Mice receiving DQF and partial antimicrobial therapy (DQF +Eth/Rif + Eth) showed a median time to death (MTD) of 98 days and displayed significantly more weight gain (by 10-15%) in comparison to those receiving partial antimicrobial therapy alone (Eth/Rif + Eth, MTD of 68 days, *p*= 0.05) (Figure 5C, 5D). As has been noted by our group and others,^34,38,40,70^ *B6.Sst1S* mice infected with this *Mtb* challenge dose display a bimodal mortality curve irrespective of therapeutic intervention, with 40-50% of all animals succumbing over the first 35 days, and thus the impact of our drug regimens was only apparent in the remaining 50-60% of surviving mice.

**Figure 5.**
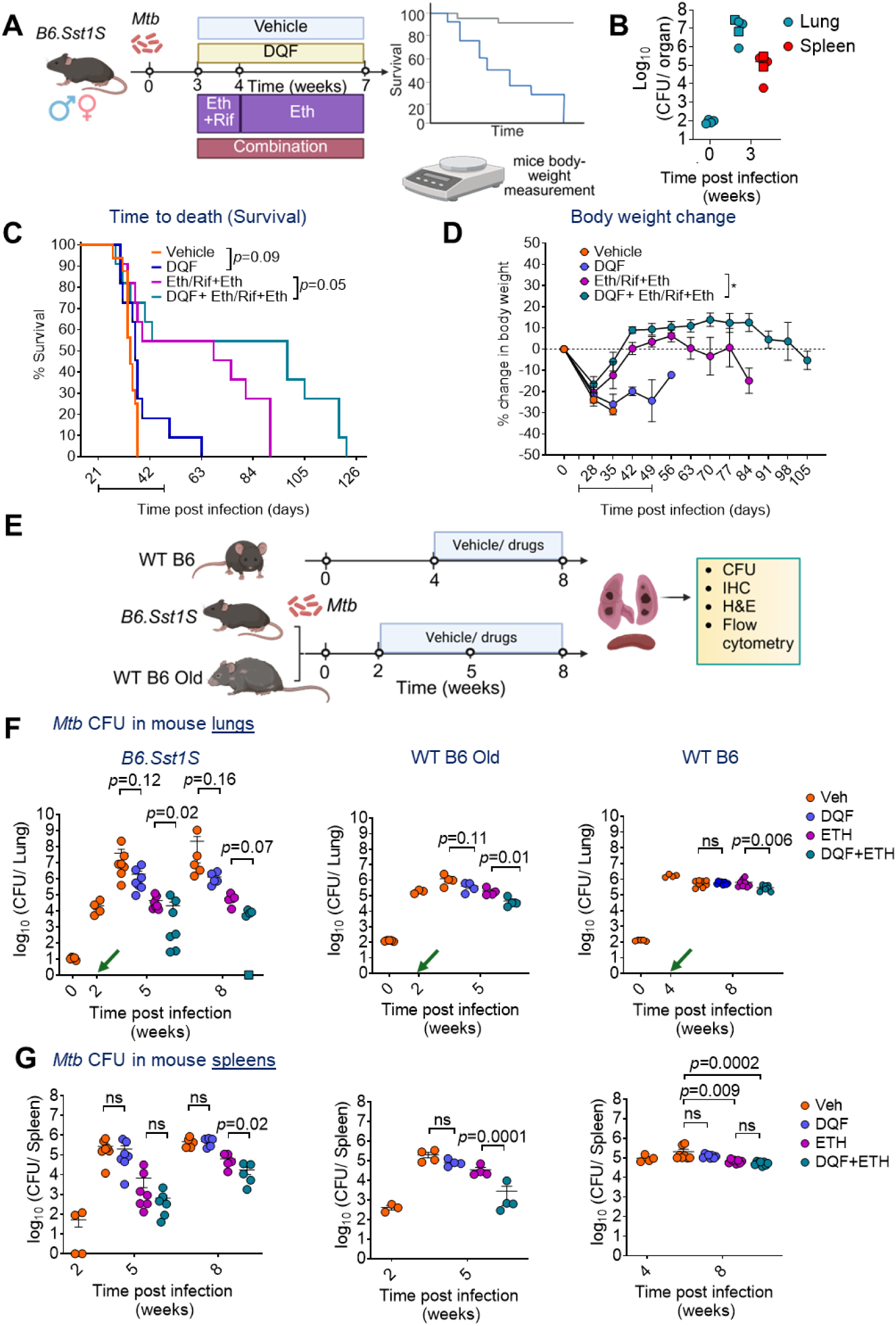
Senolytics DQF administration with or without anti-TB drug improved survival, body weight and reduced *Mtb* burden in mice. (A) Both male and female *B6.Sst1S* mice were aerosol infected with high-dose 91 CFU of *Mtb* H37Rv and at 3 weeks p.i., mice were oral gavaged with vehicle (n=16 mice)/ drugs (n=11 mice) for the next 4 weeks. (B) At 3 weeks p.i., the *Mtb* burden in lungs and spleen were 1.5*10^7^ and 1.6*10^5^ CFU, respectively. (circles indicate females and squares indicate males, n=4-5 mice). Mice were monitored for changes in body weight and survival after the start of drug treatment. (C) Kaplan-Meir survival curve of *B6.Sst1S* mice. Line indicates duration of therapy. Indicated *p* values are measured by log-rank (Mantel-Cox) test. (D) Changes in body weight are plotted as a % change in body weight of surviving mice relative to Day 1 p.i. over time. The data are means ± SEM. (*p*<0.05: *, two-way ANOVA with Tukey’s multiple comparisons test). (E) The experimental strategy. Day 1 CFU in *B6.Sst1S* mice= 11 (low dose); WT B6 young and old mice= 126. *Mtb* H37Rv burden in (F) lung and (G) spleen of WT B6, *B6.Sst1S* and old WT B6 mice at indicated time points (n=4-7). Square indicates CFU below the limit of detection (=10 CFU). The green arrow indicates the start of drug treatment. The data are means ± SEM. Each data point represents a mouse. n= 4-7 mice/ group. Statistical analysis was done by one-way ANOVA with Sidak’s multiple comparison test or Kruskal-Wallis Dunn’s multiple comparison test (5G-WT B6).

### The DQF senolytic drug cocktail reduces *Mtb* proliferation and improves lung pathology in young *B6.Sst1S* and aged WT B6 mice

Next, we evaluated the anti-TB effect of the DQF cocktail with or without Eth-antimicrobial therapy in a lung CFU burden study of young *B6.Sst1S*, aged WT B6, and young WT B6 mice, following a low-dose *Mtb* aerosol challenge. (A low challenge dose was selected to ensure that vehicle-treated *B6.Sst1S* mice would survive until the experimental endpoint). In this experiment we started drug treatment at a later time point for WT B6 mice (4 wpi) than for young *B6.Sst1S* and aged WT B6 (2 wpi) as shown in Figure 5E based upon our prior observations that WT B6 mice showed lower levels of senescence markers at early timepoints and were better able to contain *Mtb* proliferation than the other two groups. Compared to the vehicle-treated group, DQF therapy reduced lung CFU counts in *B6.Sst1S* mouse lungs by 1.3 (*p*= 0.12) and 2.2 (*p*= 0.16) log_10_CFU units at 5 and 8 wpi, respectively, and in aged WT B6 mouse lungs by 0.5 (*p*= 0.11) log_10_ units at 5 wpi (Figure 5F), but none of these reductions were statistically significant. We did not find a detectable difference in lung CFU counts of young WT B6 mice at 8 wpi (Figure 5F). The addition of ethambutol to DQF host-directed therapy significantly lowered lung CFU counts in all mouse strains compared to ethambutol monotherapy; however, the magnitude of the reduction was most profound in *B6.Sst1S* mice at 0.31(*p*= 0.02) and 1.0 (*p*=0.07) log_10_CFU units at 5 and 8 wpi, respectively, and in aged WT B6 mice at 0.6 log_10_CFU (*p*= 0.01) units at 5 wpi (Figure 5F). The difference in treatment times between mice strains precluded us from comparing outcomes across groups. However, in the given experimental conditions, young WT B6 mice also showed a reduction of 0.3 log_10_CFU units at 8 wpi (*p*=0.006) upon adding DQF to ethambutol (Figure 5F). With spleen CFU counts, the addition of DQF to ethambutol lowered the bacterial burden in *B6.Sst1S* mice by 1.0 and 0.6 log_10_CFU units at 5 (*p*=0.24) and 8 wpi (*p*=0.02), respectively. In case of aged WT B6 mice, spleen CFU burden was reduced by 1.1 log_10_CFU units at 5 wpi (*p*= 0.0001) compared to ethambutol alone (Figure 5G). There was no significant reduction observed between these groups for young WT B6 mice (Figure 5G).

Next, we examined whether lung immunopathology is altered by senolytic drug therapy in the above experimental setup. In line with previous studies, necrotic granulomas were observed in 3 out of 6 (50%) *B6.Sst1S* mice at 5 wpi, and 3 out of 5 (60%) mice at 8 wpi (Figure S8A, S8B). Treatment with DQF alone delayed the emergence of necrotic granuloma formation, as necrotic granuloma was observed in 1 out of 6 mice (17%) and in none of the 5 mice (0%) at 5 and 8 wpi, respectively (Figure S8A, S8B). Additionally, treatment with bacteriostatic drug Eth alone also resolved lung pathology and none of the mice exhibited necrosis (Figure S8A, S8B). Next, we measured the percentage area (of the lung) of consolidated lesions on H&E-stained histology slides (Figure 6A). All three mouse strains showed greatest reductions in percent lesion area in the DQF+ Eth-treatment group as compared with the vehicle group (*B6.Sst1S*: *p*= 0.02, aged WT B6: *p*= 0.07, and young WT B6: *p*= 0.04) (Figure 6A). While DQF alone prevented the formation of necrotic granuloma in *B6.Sst1S* mice, no statistically significant difference in percent lesion area was observed (Figure 6A). Additionally, the overall degree of lung consolidation remaining (percent lesion area) was improved in DQF+ Eth-treated *B6.Sst1S* (5 wpi-≈5-fold, 8 wpi-≈4-fold) and aged WT B6 mice (5 wpi-≈2-fold) relative to the vehicle-treated mice (Figure 6A, Figure S8, S9A). In the case of young WT B6 mice, Eth alone was insufficient to improve the lung pathology at 8 wpi. However, DQF alone (22%; *p*= 0.001) and DQF+Eth (26%; *p*= 0.04) treatment significantly reduced the percentage lung lesion area relative to the Veh group (37%) (Figure 6A).

**Figure 6.**
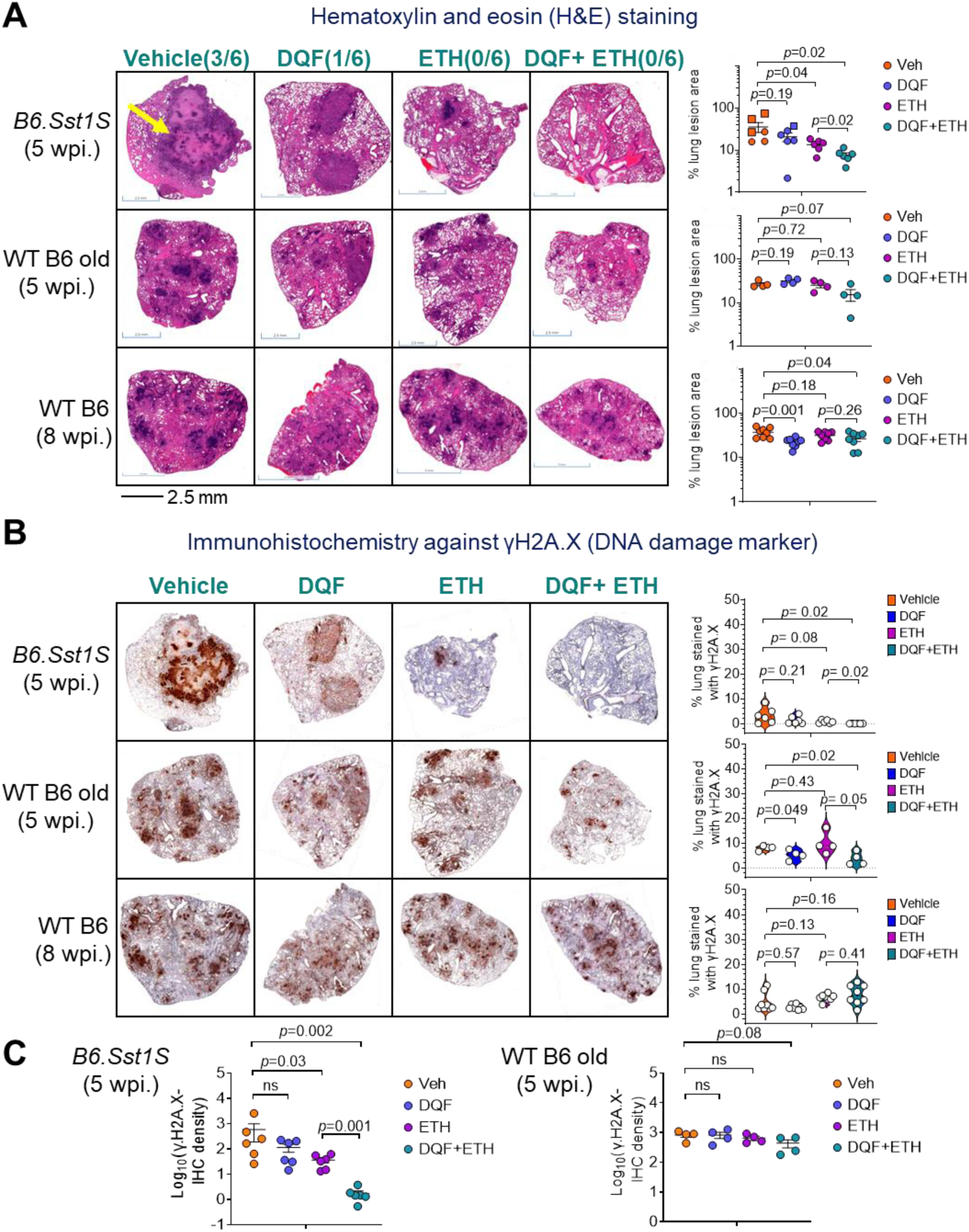
Senolytics administration alone, and in combination with Ethambutol reduced lung pathology and senescence-associated marker in *Mtb* H37Rv infected *B6.Sst1S* and WT B6 old mice. (A) Representative H&E-stained images of *Mtb*-infected mice lungs treated with Vehicle or drugs, and respective ImageJ-based quantification. Yellow arrow indicates necrotic granuloma in Vehicle treated *B6.Sst1S* mice at 5 wpi. The number in parenthesis indicate the number of mice with necrotic granuloma at 5 wpi. (B) Representative γH2A.X-Immunohistochemistry images of mice after lungs vehicle/ drugs treatment, and respective Violin plots to show ImageJ quantification of γH2A.X-stained area and (C) γH2A.X-stain intensity at indicated time points. Each data point represents a mouse. n=4-7 mice/group. Statistical analysis between two groups was done by unpaired two-tailed Student’s t test or Mann-Whitney test.

In sum, CFU and histology data suggest that the DQF senolytic drug cocktail can act as a valuable host-directed therapy in both young *B6.Sst1S* and aged WT B6 mouse TB models.

### The DQF cocktail reduces senescence markers in *Mtb*-infected lungs of young *B6.Sst1S* and aged WT B6 mice

To determine if the improved survival, *Mtb* containment, and lung pathology observed in DQF-treated mice correlated with the removal of senescent cells, we evaluated markers of senescence (γH2A.x, p21, p16 and SA-βGal activity) in mouse lungs for the 3 categories of mice treated as in Figure 5E. Immunohistochemistry for the DNA damage marker γH2A.X showed no significant differences in immunoreactivity across mice strains between DQF-alone treated and vehicle-treated mice, but in aged WT mice (*p*=0.049) (Figure 6B, Figure S9B). Interestingly, Eth treatment reduced DNA damage area of the lung by 4-fold only in *Mtb*-infected *B6.Sst1S* mice at 5 wpi (Figure 6B); however, this effect was not seen at 8 wpi (Figure S9B). Moreover, Eth-treated young or aged WT B6 mice exhibited similar γH2A.X-immunoreactivity relative to the vehicle group (Figure 6B). Importantly, the highest reduction of DNA damage was observed in DQF+ Eth-treated *B6.Sst1S* (≈80-fold at 5 wpi relative to the vehicle group) and aged WT B6 mice (≈2-fold at 5 wpi relative to the vehicle group) with mice receiving both drugs showed a near absence of any γH2A.X-stained lung area (Figure 6B, Figure S9B). In terms of the intensity of γH2A.X-staining, the highest reduction of DNA damage was also observed in DQF+ Eth-treated *B6.Sst1S* with a ≈350-fold reduction at 5 wpi relative to the vehicle group, and ≈4-fold lower for aged WT B6 mice at 5 wpi (Figure 6C).

Notably, when we evaluated the live lung cells by flow cytometry for dual staining with p21^+^βGal^+^ and for p16^+^βGal^+^ (Figure S4), there were significant reductions in both pairs of senescence markers in the DQF alone group and in the DQF+ Eth group for young *B6.Sst1S* and aged WT B6 mice (Figure 7A, 7B Figure S9C, S9D). Eth monotherapy reduced p21^+^βGal^+^ and p16^+^βGal^+^ population by ≈10-fold in *B6.Sst1S* mice lung at 5 wpi, indicating that inhibiting *Mtb* replication can also impede the generation of p21+βGal+ or for p16+βGal+ cells (Figure 7A, 7B). Interestingly, DQF alone also reduced the p21^+^βGal^+^ and for p16^+^βGal^+^ by ≈4-fold relative to the vehicle group in *B6.Sst1S* mice, and we did not see further reduction of p21^+^βGal^+^/p16^+^βGal^+^ population when DQF was added to the Eth monotherapy (Figure 7A). In the case of aged WT B6 mice, while Eth monotherapy was ineffective (*p*>0.05), DQF+ Eth significantly reduced the senescence markers in the mice lung (Figure 7B). In contrast, we saw no changes for dual positivity in young WT B6 mice with any of the DQF and/or Eth regimens compared to vehicle-treated group (Figure 7C). The lung cells showing the highest reductions in dual positivity upon DQF or Eth treatment for p21+βGal+ or for p16+βGal+ were predominantly from the CD45+myeloid lineage cells (Figure 7D, 7E).

**Figure 7.**
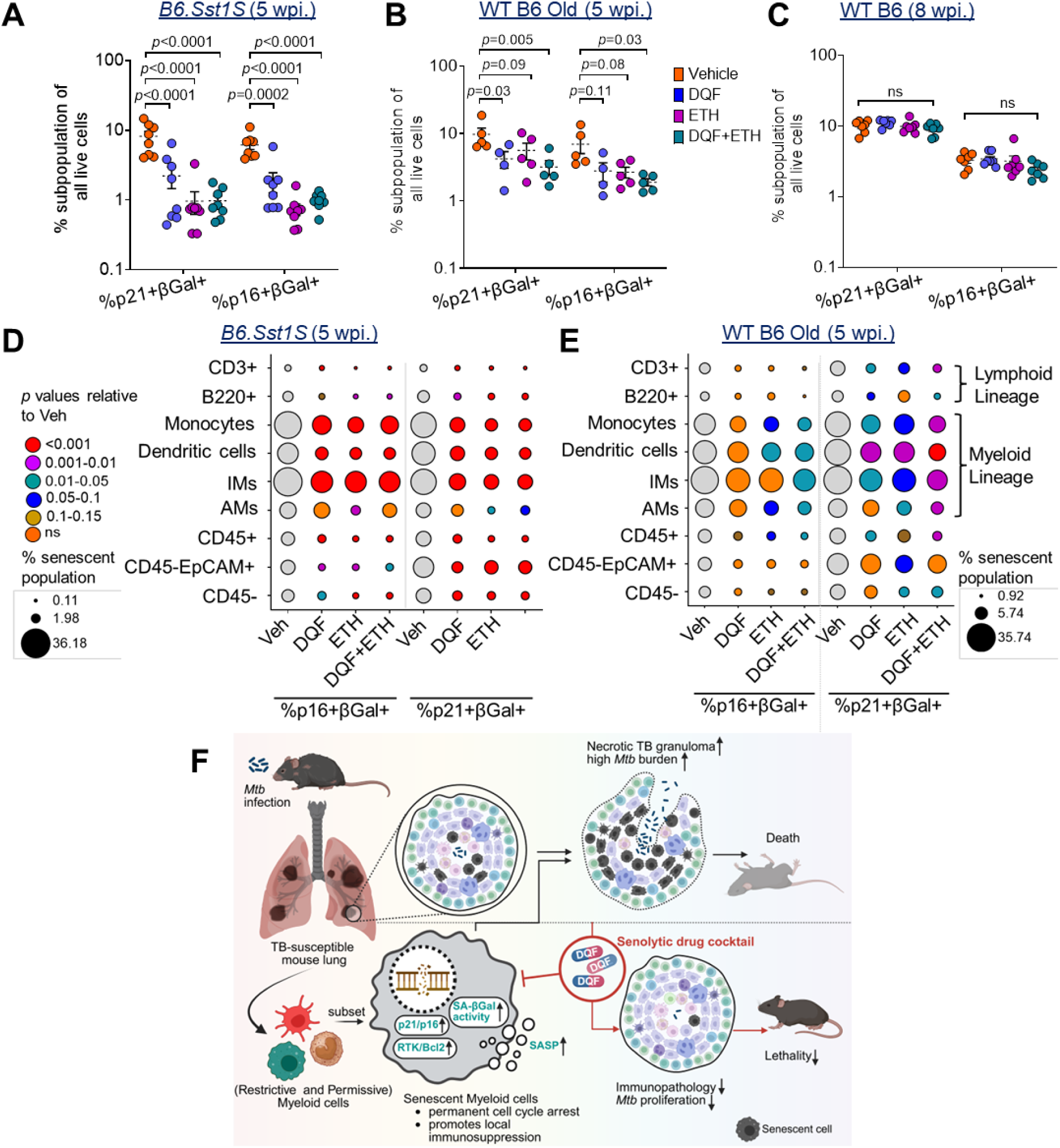
Senolytics administration alone, and in combination with Ethambutol reduced p21+SA-βGal+ or p16+SA-βGal+ senescent population in *Mtb* H37Rv infected *B6.Sst1S* and WT B6 old mice. %p21+βGal+ and %p16+βGal+ subpopulation observed in all live lung cells of Mtb-infected (A) *B6.Sst1S* (B) WT B6 old, and (C) young WT B6 mice (n= 5-8),. The data are means ± SEM. Each data point represents a mouse. Statistical analysis was calculated by two-way ANOVA with Dunnett’s multiple comparisons test. Bubble plot to show %p21+βGal+ and %p16+βGal+ subpopulation in different lung cell types in *Mtb*-infected (D) *B6.Sst1S* mice (n= 8) and (E) WT B6 old mice (n= 5) at indicated timepoint and treatment groups. The size of the bubble indicates %subpopulation out of all of particular cell types and color is adjusted *p* values relative to the Veh group. Statistics are calculated by one-way ANOVA with Tukey’s multiple comparisons test. (F) Proposed model showing the crosstalk between senescence and necrotic granuloma trajectory.

In summary, the senolytic drug cocktail DQF prolonged survival, lowered lung CFU counts, and pathology concomitant with a reduction in senescence markers in TB-infected mice (Figure 7F). The greatest host-beneficial effects were observed in TB-susceptible young *B6.Sst1S* and aged WT B6 mice.

## Discussion

This study provides proof of concept data that lung myeloid cells acquire a senescence phenotype during TB and that these senescent cells contribute to disease progression. In this work, we first used BMDMs to establish that TNFα treatment in aged WT B6 and young *B6.Sst1S* BMDMs are not only associated with poorer *Mtb* containment but also the induction of classical senescence markers including SA-βGal, SASP gene expression, cell cycle arrest (reduced BrdU incorporation, Lamin B1 and increased p16), and DNA damage (γH2A.X). Our previous work with the *B6.Sst1S* macrophages revealed up-regulation of many SASP genes (*Il6, Ccl2, Cxcl11, Tgfβ3, Mmp13, Tnfsf10*) (Supplementary File S1) and induction of the anti-apoptotic ISR pathway after TNFα-stimulation at an early time point (18 hours).^38^ Using *Mtb*-infected aged WT B6 mice as a benchmark, we sought to determine whether young *B6.Sst1S* mice resembled aged WT B6 or young WT B6. Our studies revealed that both young *B6.Sst1S* mice-lungs and their BMDMs displayed the same panel of senescence markers during stress (TNFα) or *Mtb* infection as observed in aged WT B6 samples. The senescence phenotypes seen in the *B6.Sst1S* model resembled those of aged WT B6 samples to a much greater degree than those of young WT B6. These data suggest that after *Mtb* infection, *B6.Sst1S* mice exhibit accelerated myeloid cell senescence, and that this may contribute to their unique development of necrotic granulomas.

Advanced age is a well-known TB risk factor,^7,62,63^ where early mouse studies by Orme and Turner showed that *Mtb*-infected old mice have accelerated lethality post-*Mtb* challenge.^14,71,72^ The macrophages in aged mice were *Mtb*-infected in higher numbers and expressed higher levels of the IFN-I responsive genes IRGM-1 and IRF-1.^73^

Interestingly, DiNardo *et al.* recently found that active TB leads to accelerated aging in humans and in a guinea pig acute TB model using the DNA methylation clock.^74,75^ Using combined hypermethylation and gene expression clock algorithms, they found that active TB adds 12.7 years (by the Horvath DNA hypermethylation clock) or 14.4 years (by the Transcriptomic ‘RNAAgeCalc’ clock) of cellular aging compared to latent TB-infected humans (LTBI).^74^ Consistent with this, TB patients have been reported to show significantly shorter telomere length and increased mitochondrial DNA copy number (markers of senescence) relative to healthy controls in a diverse population.^76–78^ Additional studies have indeed demonstrated that *Mtb* infection induces DNA damage, manipulates host cell cycle checkpoints, and promotes senescence-like features in macrophages,^79–82^ or T cells.^83^

The senescence literature has demonstrated that pathways required for homeostatic development and wound healing in young age become dysregulated during aging.^30^ These include fibrosis with stem cell exhaustion in the setting of chronic mTOR activation, persistent inflammation with excessive SASP cytokine release due to chronic NF-κB activation, DNA damage and mitochondrial dysfunction in the setting of declining sirtuin activity.^84,85^ A hallmark of senescent cells is entry into a non-dividing, long-lived state with chronic expression of locally deleterious and immunosuppressive signals that promote age-associated disease processes. In TB, the majority of infected humans adequately contain *Mtb* infection within immune-balanced quiescent lung granulomas. We propose that in TB-susceptible hosts, such as *B6.Sst1S* mice (interferonopathy) or aged WT B6 mice (aging/inflammaging), a subset of stressed myeloid cells transit into a stress-induced senescent (SIS) state characterized by growth arrest and local secretion of SASP factors.

These SASP molecules include type I interferon, PD-L1, IL-1 receptor antagonist,^34^ IL-10, IL-6, CCL2, and GM-CSF. These senescent myeloid cells, via SASP secretion, may promote chronic local inflammation and recruit immunosuppressive myeloid-derived suppressor cells (MDSCs) into granulomas.^25,26,86^ We indeed observed several canonical immunoregulatory molecules that can collectively dampen antimicrobial immunity, to be upregulated in the cells neighboring to non-controlling lesions in *B6.Sst1S* mice (Supplementary File S2). Among the upregulated DEGs, Checkpoint receptors or ligands such as Pdcd1 (PD-1; T-cell exhaustion), Cd274 (PD-L1), Fcgr2b, Cd33, Ceacam1, Sirpb1a and Tnfrsf1b can prevent T cell activation. Metabolic and enzymatic brakes, including Arg1, Arg2, Nos2, Il4i1, Ptgs2, Acod1, Hmox1 and the adenosine receptor Adora2b, constrain lymphocyte proliferation by depleting arginine or generating immunomodulatory metabolites. Signal-terminating proteins Socs3, Nlrp12, Dusp1, and the IL-1 antagonist Il1rn attenuate JAK/STAT, NF-κB or MAPK cascades, whereas chemokines-Ccl2,Ccl3, Cxcl1, Cxcl2 and S100A8/S100A9-alarmin can recruit MDSCs,^87–89^ reinforcing a suppressive milieu locally in the tissue (Supplementary File S2). In turn, it has been shown that MDSCs further sustain immunosuppression by secreting IL-10, TGF-β, and reactive oxygen species (ROS), impairing effective cytotoxic T-cell and NK-cell responses and preventing clearance of senescent cells.^27^ These paracrine signals also feed the cycle of IFN-I hyperexpression in uninfected bystander cells, blunt T cell activation,^44^ and recruit additional immature myeloid cells leading to granuloma progression. Consequently, SIS myeloid cells and recruited MDSCs may collectively shape an immunosuppressive granuloma microenvironment favorable to *Mtb* replication, while impeding the establishment of protective immune control, as seen in cancer-biology.

We found that senescence markers in aged WT B6 and young *B6.Sst1S* appear as early as 10 days after *Mtb* infection, well before the appearance of granuloma formation and lung immunopathology. This early chronology suggests that myeloid cell senescence may be a contributory driver of the more severe TB pathology in aged WT B6 and young *B6.Sst1S* animals. To establish causality, we used a cocktail of the senolytic drugs dasatinib, quercetin, and fisetin (DQF) which eliminate senescent cells by inhibiting the kinase (RTK/ PI3K/ ERK)-dependent survival pathways and anti-apoptotic mechanisms that sustain their long-lived, non-dividing state. These drugs have been shown to prolong lifespan in rodents and to prevent the progression of age-associated diseases in humans.^21,58,60,61^ We found that the DQF cocktail with or without partial antimicrobial therapy was sufficient to prolong mouse survival and reduce *Mtb* organ burden. Moreover, the impact of the senolytic DQF cocktail was more prominent in mice that showed elevated senescence markers during *Mtb* infection, namely young *B6.Sst1S* and aged WT B6 mice. The fact that senolytic drugs improve TB disease outcome parameters is a potent argument that senescent lung myeloid cells contribute to TB progression in these animal models. As major risk factors for TB (age, malnutrition, diabetes, pollution, and smoking) have been shown to induce senescence,^72,90–95^ this study correlating senescent cells with TB progression may also offer a mechanistic link for how these human risk factors promote TB progression. Interestingly, we also observed that treatment with the bacteriostatic drug ethambutol reduced the accumulation of senescence markers. This supports the established notion that chronic lung inflammation resulting from persistent *Mtb* growth can indeed induce stress-induced senescent (SIS) cells. Taken together, our findings suggest that TB susceptibility factors promote a lung environment favorable for *Mtb* replication, fueling a self-amplifying/feed-forward cycle of senescence and inflammation, ultimately driving progression toward necrotic granulomas.

Considerable recent attention has been directed to the problem of post-TB lung disease. Despite adequate and durable cures, many TB patients are left with serious pulmonary disease, including bronchiectasis and either obstructive or restrictive lung disease.^96–99^ Senescent cells have been strongly associated with chronic tissue inflammation and inappropriate fibrotic scarring.^100^ Our studies with senolytic drugs indicated that lung immunopathology was reduced to a greater degree by senolytic drugs in young *B6.Sst1S* and aged WT B6 mice, which have high senescent cell burdens. Thus, in addition to serving as promising adjunctive host-directed therapies for active TB, senolytic drugs may also reduce post-TB lung disease. In summary, this demonstration of senolytic drug efficacy mouse TB models offer a rationale for testing senescence-targeting strategies in human TB clinical trials.

## Material and Methods

### Animals used in this study

Young C57BL/6J female mice (6 weeks old; Strain: 000664) and Aged C57BL/6J females (>68 weeks old) were bought from The Jackson Laboratory (Bar Harbor, ME, USA). The B6.C3H-sst1(B6J.C3-sst1C3HeB/FejKrmn) congenic mice were generated previously by transferring the *sst1S* allele from C3HeB/FeJ mouse strain to the B6 (C57BL/6J) genetic background using 12 backcrosses (*B6.Sst1S*)^32,38^ and confirmed by Genotyping. The oligonucleotides used for the PCR are shown in Supplementary File S6. This mice strain was bred and maintained at the Rodent Vivarium of Johns Hopkins University, School of Medicine and was used in this study. All experiments were conducted under protocols approved by the Johns Hopkins Animal Care and Use Committee (Protocol Number: MO22M466) and followed established ethical animal handling and care guidelines. Animals were housed in individually ventilated cages under controlled conditions, including a 12-hour light/dark cycle, ambient temperatures ranging from 20 to 24°C, and relative humidity between 45% and 65%. Each cage contained no more than five mice of the same strain/sex to ensure proper animal welfare. Mice were randomly assigned to experimental groups to minimize bias. Humane and painless euthanasia procedures were employed in compliance with AVMA (American Veterinary Medical Association) guidelines.

All experiments involving *Mycobacterium tuberculosis* H37Rv were conducted in Biosafety Level 3 (BSL3) and Animal Biosafety Level 3 (ABSL3) facilities at the Johns Hopkins University, School of Medicine, following protocols approved by the Institutional Biosafety Committee. Aerosol infections were performed using a Glas-Col aerosolization system, allowing the mice to move freely in the chamber and minimizing stress or discomfort. Anesthesia was not required for this procedure. Mice were briefly restrained during oral gavage to ensure accurate delivery. Infected animals were monitored for signs of illness, such as weight loss or lethargy, and those that became moribund before the experiment’s conclusion were promptly euthanized, except in survival studies.

### BMDMs isolation, culture, and treatment with murine-TNFα

BMDMs were isolated from mice at the indicated age, as described previously^101^. Briefly, BMDMs were isolated from the tibias and femurs of mice and were differentiated into macrophages in RPMI-Glutamax (Thermo Fisher Scientific; 61870127) containing 10% heat-inactivated FBS (Millipore Sigma, Catalog no.-F4135), 1X antibiotic-antimycotic Solution (Sigma Aldrich; A5955) and 10% (v/v) L929 fibroblast-conditioned medium (as a source of M-CSF(macrophage colony-stimulating factor)) for 5 days. L929 fibroblast cell lines were cultured in DMEM-Glutamax (Thermo Fisher Scientific; 10566016) containing 10% heat-inactivated FBS, 10 mM HEPES, and 1X antibiotic-antimycotic Solution. L929 cells were mycoplasma-negative (Venorgem Mycoplasma detection kit; Millipore Sigma-MP0025). BMDMs were either left untreated or treated with 10 ng/ml of murine-TNFα (Peprotech; 315-01A) in RPMI-Glutamax containing 10% heat-inactivated FBS and 10% (v/v) L929 fibroblast-conditioned medium, for indicated time durations at 37°C in 5% CO_2_. BMDMs were washed, scraped out, counted, and replated overnight prior to the experiments. All experiments were done with confluency of BMDMs 60-75%.

### Senescence associated β-Galactosidase assay (SA-βGal assay)

SA-βGal assays at indicated time points were done with both paraformaldehyde-fixed (Senescence Cells Histochemical Staining Kit; Sigma; CS0030) and live BMDMs (Senescence β-Galactosidase Activity Assay Kit (Fluorescence, Flow Cytometry); Cell Signaling Technology; 35302), using a modified manufacturer’s protocol. At indicated time points, BMDMs were washed twice with PBS and fixed with 1X fixation buffer for 10 minutes. BMDMs were washed thrice, the staining mixture was added to each well and incubated for 48 h at 37°C without CO_2._ We used 0.04X the X-gal Solution (10 µl in 1ml of Staining mixture), to avoid saturation of the signal. The plate was sealed with Parafilm to prevent evaporation of the staining mixture. After the incubation, cells were observed under a bright field microscope and images were captured from the center of the well.

For experiments with live cells, adherent BMDMs were washed twice with warm PBS, and incubated in Bafilomycin A1 (freshly prepared; 100 nM) containing complete RPMI medium for 2 hours. Next, 0.05X of the recommended concentration of the SA-βGal Substrate Solution (0.66 µM) was added to the cells and incubated for another 2-2.5 hours at 37°C and 5% CO_2_. The cells were washed thrice with PBS, scraped out in 1ml RPMI (without phenol red) (Gibco; 11835), and LIVE/DEAD Fixable Near-IR Dead Cell Stain (Thermo Fisher Scientific; L10119) was added for the detection of dead cells. FACS was immediately done using CytoFLEX cytometer (Beckman Coulter), and 10000 events/sample were recorded. BMDMs from aged WT B6 mice (>72 weeks) were used as a positive control. For etoposide (Millipore Sigma; E1383)-induced senescence control, WT B6 BMDMs were treated with 10 µM etoposide for 2 days in complete RPMI media, followed by washing to remove DNA-damaging molecules, and further incubation for 2 days, in complete RPMI media only (no loss of viability under these conditions, as detected by Alamar HS assay).

### Cell viability assay

The viability of cells was measured using an Alamar Blue HS Cell Viability Reagent (Thermo Fisher Scientific; A50100), according to the manufacturer’s protocol. Briefly, at the indicated point, the (warmed) viability reagent was added to each well at 1/10^th^ the volume of cells, incubated for 6-8 hours and the fluorescence intensity was measured using a FLUOstar Omega Multidetection Microplate Reader (BMG Labtech) in the bottom-reading mode with excitation at 544 nm and emission at 590 nm. Untreated cells and cells treated with 0.06% SDS, were used as positive and negative control, respectively. Percent viability was calculated from the relative fluorescence units compared with positive and negative control cells, to which the experimental values were normalized.

### Isolation of RNA and quantitative RT-PCR analysis to detect SASP expression

At the indicated time points, BMDMs were lysed in TRIzol (Thermo Fisher Scientific; 15596026), and RNA was extracted using Qiagen RNeasy Mini Kit (74104) and RNase-free DNase Set (Qiagen; 79254), following the manufacturer’s protocol. RNA purity and concentration were quantified using Nanodrop One (Thermo Fisher Scientific). 1 µg of RNA was used for cDNA synthesis using the iScript cDNA Synthesis Kit (BioRad; 1708891) and quantitative RT-PCR was done with QuantStudio 3 (Applied Biosystems; Thermo Fisher Scientific) using gene-specific primers (Supplementary File S6) and iTaq Universal SYBR Green Supermix (BioRad; 1725121). Expression of genes was normalized with Ct value for *ActB* (internal housekeeping control).

### BrdU Cell proliferation assay

The capacity of BMDMs to proliferate was measured by using the BrdU Cell Proliferation Assay Kit (Cell Signaling Technology; 6813), following the manufacturer’s protocol. Briefly, 1-day post seeding of 50000 BMDMs in a 48-well plate, growth media (RPMI-Glutamax+ FBS) was removed, and RPMI-only was added to the cells to synchronize their cell cycle (upon serum starvation) and incubated for 18 hours at 37°C and 5% CO_2_. After this synchronization-step, complete RPMI media was added to all wells and incubated for another 24 hours. The cells were either left untreated or treated with 10ng/ml TNFα, and BrdU incorporation into the cells was measured following the manufacturer’s recommendations. Treatment of WT B6 BMDMs with 10 µM etoposide acted as a positive control. Absorbance at 450 nm was measured using the Bio-Rad iMark microplate reader.

### Protein isolation and Western Blotting

BMDMs were washed twice with ice-cold PBS, and the total protein was extracted by lysing cells in radioimmunoprecipitation (RIPA) Buffer (Millipore Sigma; R0278) containing Protease/Phosphatase Inhibitor Cocktail (1X; Cell Signaling Technology; 5872). After incubating on ice for 15 min with gentle vortexing, the lysates were centrifuged at 14000 RPM, 4°C for 15 min. The clarified supernatant was taken, and protein concentrations were quantified using the BCA Assay kit (Thermo Fisher Scientific; 23225). Approximately 50 μg of protein were mixed with Laemmli sample buffer (Bio-Rad; 1610747) and heated at 95 °C for 5 min before loading onto 4-15% MP TGX polyacrylamide Gel (Bio-Rad; 45610848) and transferring to the PVDF membranes (Bio-Rad; 1620177). The PVDF membranes were blocked using 5% Blotting-grade blocker (Bio-Rad; 1706404) in TBST buffer-Tris-buffered saline (Quality Biological; 351086101)+ 0.1% Tween 20 (Sigma; P2287) for 1-2 hour and incubated overnight at 4°C with the primary antibodies from Mouse Reactive Senescence Marker Antibody Sampler Kit (Cell Signaling Technology-78551)-Phospho-Histone H2A.X (Ser139) (20E3) Rabbit mAb (1: 1000 dilution), Lamin B1 (E6M5T) Rabbit mAb (1: 1000 dilution), HMGB1 (D3E5) Rabbit mAb (1: 1000 dilution), p16 INK4A (E5F3Y) Rabbit mAb (1: 1000 dilution) and β-Actin (D6A8) Rabbit mAb (Cell Signaling Technology; 8457) (1:5000 dilution). Following three 15 min washing steps with TBST, the membranes were incubated with secondary antibody-Anti-rabbit IgG, HRP-linked Antibody (1:1000 dilution) for 2 hours at room temperature (RT). Following three 15 min washing steps, the membranes were developed with Pierce™ ECL Western Blotting Substrate (Thermo Fisher Scientific; 32109) and visualized using a KwikQuant Digital Western Blot Detection System (Kindle Bioscience; D1001). Images were processed in Adobe Photoshop Software. Stripping Buffer (Restore™ Western Blot Stripping Buffer; Thermo Fisher Scientific; 21059) was used to redevelop PVDF membranes, following the manufacturer’s protocol.

### Bulk RNA-Sequencing and analysis

BMDMs were lysed in TRIzol (Thermo Fisher Scientific; 15596026), and RNA was extracted using Qiagen RNeasy Mini Kit (74104) and RNase-free DNase Set (Qiagen; 79254), following the manufacturer’s protocol. RNA purity and concentration was quantified using Nanodrop One (Thermo Fisher Scientific) and stored at −80 °C. RNA samples were shipped on dry ice to Genewiz, Azenta Life Sciences (South Plainfield, NJ, USA) for standard bulk RNA-sequencing and downstream analysis. Sample QC, library preparations, sequencing reactions, and initial bioinformatic analysis were conducted as follows: Total RNA samples were quantified using Qubit 2.0 Fluorometer (Life Technologies, Carlsbad, CA, USA) and RNA integrity was checked with 4200 TapeStation (Agilent Technologies, Palo Alto, CA, USA). Samples were initially treated with TURBO DNase (Thermo Fisher Scientific, Waltham, MA, USA) to remove DNA contaminants. The next steps included performing rRNA depletion using QIAseq® FastSelect™−rRNA HMR kit (Qiagen, Germantown, MD, USA), which was conducted following the manufacturer’s protocol. RNA sequencing libraries were generated with the NEBNext Ultra II RNA Library Preparation Kit for Illumina by following the manufacturer’s recommendations. Briefly, enriched RNAs were fragmented for 15 minutes at 94°C. First strand and second strand cDNA were subsequently synthesized. cDNA fragments were end-repaired and adenylated at 3’ends, and universal adapters were ligated to cDNA fragments, followed by index addition and library enrichment with limited cycle PCR. Sequencing libraries were validated using the Agilent Tapestation 4200 (Agilent Technologies, Palo Alto, CA, USA), and quantified using Qubit 2.0 Fluorometer (ThermoFisher Scientific, Waltham, MA, USA) as well as by quantitative PCR (KAPA Biosystems, Wilmington, MA, USA). The sequencing libraries were multiplexed and clustered on one lane of a flow cell. After clustering, the flow cell was loaded on the Illumina HiSeq 4000 instrument according to the manufacturer’s instructions. The samples were sequenced using a 2×150 Pair-End (PE) configuration. Image analysis and base calling were conducted by HiSeq Control Software (HCS). Raw sequence data (.bcl files) generated from Illumina HiSeq was converted into FASTQ files and de-multiplexed using Illumina’s bcl2fastq 2.17 software. One mismatch was allowed for index sequence identification.

Raw sequencing reads were trimmed to remove adapter sequences and low-quality bases using Trimmomatic v0.36, and the filtered reads were aligned to the *Mus musculus* GRCm38 reference genome (ENSEMBL) using the splice-aware STAR aligner v2.5.2b, generating BAM files. Gene hit counts were quantified using featureCounts from the Subread package v1.5.2, counting only uniquely mapped reads overlapping annotated exon regions, with strand specificity applied where appropriate. Differential expression analysis was performed using the DESeq2 package^102^. Genes with an adjusted *p*-value <0.05 and a log₂ fold change > 1 or <-1 were deemed differentially expressed between groups. Statistically significant differentially expressed genes (DEGs) were analyzed for biological process enrichment through gene ontology analysis using the software GeneSCF v1.1-p2 with the MGI GO database, generating clusters of functionally relevant GO terms. Raw data were deposited in NCBI’s Gene Expression Omnibus (GEO, accession number-GSE292458, GSE301446). Reviewers can access this data using the following token number-ihyhoqmkbhwxxmp, cdybcgoyrbqhtap, respectively.

### Minimum inhibitory concentration (MIC) and Minimum bactericidal concentration (MBC) determination against *Mycobacterium tuberculosis* H37Rv

*Mycobacterium tuberculosis* strain H37Rv (*Mtb*) was obtained from the Johns Hopkins University, Center for Tuberculosis Research (CTBR) and sequence-verified regularly. *Mtb* was grown in 7H9 broth supplemented with 0.2% glycerol, 0.05% Tyloxapol (Millipore Sigma; T0307), and OADC (BD; 212351) with shaking at 180 rpm in a rotary shaker incubator or on 7H11 agar supplemented with OADC at 37°C. MIC was determined by a resazurin microtiter assay (REMA) using 96-well flat-bottom plates^103^. *Mtb* H37Rv was cultured in 7H9+OADC medium and grown to an exponential phase (optical density at 600 nm = 0.6-0.8). Approximately 1 × 10^5^ bacteria per well were added in a total volume of 200 μL of 7H9+OADC medium. Wells lacking *Mtb* served as controls. Additional controls consisted of wells containing *Mtb* cells without drug treatment (growth control). After 5 days of incubation at 37°C in the presence of drugs, 30 μL of 0.02% resazurin (Sigma-Aldrich; R7017) was added, and plates were incubated for an additional 24-48 hours. The fluorescence intensity was measured using a FLUOstar Omega Multidetection Microplate Reader (BMG Labtech) in the bottom-reading mode with excitation at 544 nm and emission at 590 nm. Percent inhibition was calculated from the relative fluorescence units compared with an untreated control culture; the MIC was taken as the lowest drug concentration that resulted in at least 90% reduction in fluorescence compared to the untreated growth control. To determine the MBC, 20 µl of cells from the respective MIC dilutions (from the 96-well plate) were added onto the 7H11+OADC agar plate. Following 8-10 days of incubation at 37 °C, the colonies were checked for *Mtb* growth. Three wells each from media containing no inoculum and, media containing *Mtb* but no drug, were used as blank and growth controls, respectively. MBC was defined as the lowest drug concentration at which no *Mtb* colonies appeared on the 7H11 plates.

### Spatial Transcriptomics of Lung from *Mtb* infected *B6.Sst1S* mice

To characterize gene expression in controlled vs uncontrolled lesions, we performed spatial transcriptomics analysis using the Nanostring GeoMX Digital Spatial Profiler (DSP) system (Nanostring, Seattle, WA).^104,105^ The lung sections we processed for spatial transcriptomics were obtained from *B6.Sst1S* mice. We used 10^6^ CFU of *Mtb* H37Rv for infection, administered subcutaneously in the hock. The samples were collected at 14 weeks post-infection. Based on routine hematoxylin and eosin histopathology, we selected lungs from 2 mice with paucibacillary controlled lesions and 2 mice with advanced uncontrolled multibacillary lesions with necrotic areas. Additional FFPE sections were stained with fluorescent CD45-, pan-keratin-specific antibodies and DAPI. Diseased regions of interest (ROI) were selected to focus on myeloid-rich areas avoiding necrosis and tertiary lymphoid tissue. Eight ROI each of controlled and uncontrolled lesions (respectively) were studied. The profiling used the Mouse Whole Transcriptome Atlas (WTA) panel which targets ∼21,000+ transcripts for mouse protein coding genes plus ERCC ( External RNA Controls Consortium) negative controls to profile the whole transcriptome, excluding uninformative high expressing targets such as ribosomal subunits.

Samples from each ROI were packaged into library for sequencing (NextSeq550, Illumina) following the procedure recommended by Nanostring. After sequencing, the data analysis workflow began with QC evaluation of each ROI based on thresholds for number of raw and aligned reads, sequencing saturation, negative control probe means, and number of nuclei and surface area. Background correction is performed using subtraction of the mean of negative probe counts. Q3 normalization (recommended by Nanostring) results in scaling of all ROIs to the same value for their 3rd quartile value. Statistical analysis was then performed to identify differentially expressed genes, using an unpaired t-test with Benjamini-Hochberg adjustment for *p* values. Raw data were deposited in NCBI’s Gene Expression Omnibus (GEO, accession number-GSE292392). Reviewers can access this data using the following token number-clozaycmjjivduj.

### Animal infection studies

Mice of the specified age and sex were infected via aerosol exposure using the Glas-Col Inhalation Exposure System (Terre Haute, IN, USA). Fresh cryo-aliquots of *Mtb* H37Rv were used for each infection, diluted in sterile 7H9 broth to empirically determined concentrations to achieve the desired bacterial load on day 1 post-infection. To minimize variability between groups, mice from all experimental groups were infected concurrently and randomly allocated to experimental arms. The bacterial implantation in the lungs was assessed one day after infection by sacrificing 3–5 mice per group and determining colony-forming units (CFUs). Mice were euthanized at specified time points (as outlined in the Results section), and the lungs and spleen were aseptically harvested, weighed and either homogenized by bead-beating for bacterial quantification (Right lung) or stored in 10% neutral-buffered formalin for histopathological analysis (Left lung). CFU enumeration was performed by plating serial dilutions on 7H11 agar supplemented with OADC, followed by incubation at 37°C for 3 to 4 weeks. Bacterial counts were normalized to the weight of the total organ, as previously described^106^.

### Histopathology to estimate lung inflammation

For histology, left lungs were fixed by immersion in 10% neutral-buffered formalin (Thermo Fisher Scientific; 5705) for at least 2 days, before it was submitted to the SKCCC Oncology Tissue and Imaging Service Core Laboratory (Johns Hopkins University, School of Medicine) for downstream processing. The lungs were paraffin-embedded, 4 µm-sectioned, and stained with hematoxylin and eosin (H&E). Slides were digitally scanned at 40× using a Hamamatsu Nanozoomer S210 digital slide scanner (Hamamatsu Photonics, Shizuoka, Japan) at 40X magnification (0.23 microns/pixel). Image files were submitted in https://digital.pathology.johnshopkins.edu/ using Concentriq LS platform (Proscia, PA, USA). Lung inflammation was quantified as a percentage of total lung area using an in-house protocol developed with ImageJ software version 1.54m (NIH, USA) to minimize human bias, through standardized software-based settings. Images were first opened in ImageJ, and the scale was set by drawing a line over the scale bar using the line tool and defining the known distance in millimeters under Analyze> Set Scale. Images were then cropped to exclude the ruler (Image > Crop) and saved. For area quantification, the Image> Adjust> Color Threshold tool was used, with consistent settings for hue (0– 255), saturation (0–255), and brightness (set based on analysis type)-full lung area=(0-250) and inflamed area= (0-160). After selecting the ROI, the threshold window was closed, and images were converted to binary format using Process> Binary> Make Binary. The Signal to Noise ratio was improved by removing the Outliers-using Process> Noise> Remove Outliers, adjusting the radius (=4) and the Threshold (=50). Measurements were extracted (Analyze> Measure), with the initial setup requiring the selection of “area” and “limit to threshold” under Set Measurements. Processed images were saved, and the steps were repeated for all images with identical parameters to ensure consistency. ROIs were then divided by the total lung area per slide to calculate the ratio of inflamed to lung tissue-area.

### Multicolor Flow cytometry to quantify senescent cell population in *Mtb*-infected mice lungs and spleen

To assess markers of senescence (p16+, p21+, and increased SA-βGalactosidase activity) in lung and spleen tissues from *Mtb*-infected mice, organs were harvested at indicated time points and dissociated using the Lung Dissociation Kit (Miltenyi Biotec;130-095-927), and the gentleMACS Octo Dissociator (program m_lung_02), according to the manufacturer’s protocol. Single-cell suspensions were obtained by filtering dissociated tissue through 70 µm mesh strainers (Corning; 431751), washing with 5 mL complete RPMI media, and centrifuging at 300g for 5 min at RT. Red blood cells were lysed using ACK lysis buffer (Quality Biological; 118156101), neutralized with complete RPMI, and centrifuged. Cells were stained for viability using a live-dead near-IR dye (1:1000 dilution in PBS; Thermo Fisher Scientific; L10119) and blocked with Rat Anti-Mouse CD16/CD32 Fc Block (Clone 2.4G2, BD Pharmingen; 553142). The panel of antibodies to detect the senescent cell population (SenPanel) was designed using FluoroFinder (https://fluorofinder.com/). For extracellular staining, cells were incubated with 50 µL of an antibody cocktail in Brilliant Stain Buffer (BD Biosciences; 563794) containing anti-mouse CD45 (Brilliant Violet 480; BD Biosciences;566095), CD11b (Brilliant Violet 650; BioLegend; 101239), CD45R/B220 (Brilliant Violet 421; BioLegend; 103239), CD3 (StarBright Violet 610; BioRad-MCA500SBV610), EpCAM (PerCP-Cy5.5; BioLegend; 118220), F4/80 (PE-Dazzle 594; BioLegend; 123145), CD11c (PE-Fire 810; BioLegend; 161105), and Siglec-F (Alexa Fluor 700; BioLegend; 155533) for 30 minutes at 4°C in the dark. After extracellular staining, cells were fixed with 4% formaldehyde, washed with PBS, and incubated with the CellEvent™ Senescence Green Flow Cytometry Assay Kit (Thermo Fisher Scientific; C10841; 1:250 dilution) in the senescence reaction buffer for 4 hours at 37°C, protected from light and without CO₂. Following incubation, cells were washed and permeabilized using Perm/Wash buffer (BD Biosciences; 554723). Intracellular staining was performed by incubating cells with a cocktail containing p14ARF/CDKN2A Antibody (PE; Novus Biologicals; NB200-111PE), p21/CIP1/CDKN1A Antibody (HJ21) (APC; Novus Biologicals; NBP2-47896APC), and CD68 (Brilliant Violet 785; BioLegend; 137035) for 60 minutes at 4°C in the dark. After staining, cells were washed, filtered through strainer-capped BD FACS tubes (BD; 352235), and analyzed via flow cytometry. 1×10⁶ events per sample were acquired using a BD LSR Fortessa™ Cell Analyzer (BD Biosciences), and the data were analyzed with FlowJo software version 10.8.1 (BD Biosciences). Single stain controls and compensation matrices were prepared using UltraComp eBeads Plus Compensation Beads (Thermo Fisher Scientific; 01-3333-41) and cells, as required. All flow antibodies were titrated to determine optimal staining concentrations, and only manufacturer-validated antibodies with demonstrated specificity were used. For the analysis, cells were first gated on singlets (FSC-H vs. FSC-A) and live cells (live-dead near-IR negative) before further selection of cell types. T cells are CD45+CD3+ and B cells are CD45+B220+. Myeloid cells: Tissue-resident Macrophages are CD45+CD68+F4/80+CD11B-CD11C+, Interstitial macrophages are CD45+CD68+ F4/80+CD11B+CD11C+SiglecF-, Monocytes are CD45+CD68+F4/80+CD11B+CD11C-, and Dendritic cells are CD45+CD68+F4/80-CD11B+CD11C+. Gating for p16⁺, p21⁺, and SA-βGal⁺ events for each cell type were established using reference cells from *Mtb*-infected WT B6 mice (set at 1 %).

### Immunohistochemistry against p21 and γH2A.X (phosphor S139) in mice lungs

Immunostaining was conducted at the SKCCC Oncology Tissue and Imaging Service Core Laboratory, Johns Hopkins University, School of Medicine. Immunolabeling for γH2A.X (phospho S139) and p21 was performed on 4 µm-thick formalin-fixed, paraffin-embedded (FFPE) tissue sections using the Ventana Discovery Ultra autostainer (Roche Diagnostics). Sections underwent onboard dewaxing and rehydration, followed by epitope retrieval in Ventana Ultra CC1 buffer (Roche Diagnostics; 6414575001) at 96°C for 64 minutes. Primary antibodies-anti-gamma H2A.X (phospho S139) [EP854(2)Y] (Abcam; ab81299; 1:1000 dilution) and Recombinant Anti-p21 antibody [EPR18021] (Abcam; ab188224; 1:1000 dilution) were applied as indicated, at 36°C for 60 minutes.

Detection of the primary antibodies was achieved using the anti-rabbit HQ detection system (Roche Diagnostics; Catalog 7017936001 and 7017812001), followed by visualization with the Chromomap DAB IHC detection kit (Roche Diagnostics; 5266645001). Sections were counterstained with Mayer’s hematoxylin, dehydrated, and mounted. Whole-slide imaging was performed at the Oncology Tissue Services Core of Johns Hopkins University using a Hamamatsu Nanozoomer S210 digital slide scanner (Hamamatsu Photonics, Shizuoka, Japan) at 40X magnification (0.23 microns/pixel). Images were visualized using the Concentriq digital pathology platform (Proscia, Philadelphia, PA) and analyzed in ImageJ following a previously established in-house protocol. Area quantification was performed using the Image> Adjust> Color Threshold tool in ImageJ, applying consistent hue (0–255), saturation (0–255), and brightness thresholds based on analysis type: full lung area (0–250) and DAB+ area (0–120). Identical thresholds and detection settings were applied for all slides to ensure consistency across analyses.

### Senescence-associated secretory phenotype (SASP)-cytokine quantification in *Mtb*-infected mice lungs

To estimate SASP-cytokine levels, lung homogenates were centrifuged at 14,000 RPM for 10 minutes at 4°C, and the resulting supernatants were filtered by a 0.2 μm filter, prior to removal from the BSL3 facility. Supernatants were flash-frozen in liquid nitrogen and stored at –80°C until cytokine quantification analysis. Cytokine levels in the lung homogenate were measured using the LEGENDplex Mouse Inflammation Panel 13-plex kit (BioLegend; 740446) following the manufacturer’s protocol. Data acquisition was performed on the CytoFLEX Flow Cytometer (Beckman Coulter), and cytokine concentrations were analyzed using the LEGENDplex data analysis software Suite (BioLegend). The amount of SASP cytokines were normalized to the protein concentration of the homogenate (as measured by the BCA Assay kit).

### Drug preparation and study arms to test Senolytics as host-directed therapy *in vivo*

Dasatinib (Cayman Chemical; 11498), Fisetin (Cayman Chemical;15246), and Quercetin (Millipore Sigma; Q4951) at required concentrations were first dissolved in 10ml DMSO (Millipore Sigma; D2650) to make a stock solution. This mixture was then added to 190 ml of the Vehicle (60% Phosal 50 PG (Lipoid LLC; 368315), 30% PEG 400 (Sigma Aldrich; 202398), and 10% Ethanol) to prepare the final Senolytics reagent. Ethambutol (Millipore Sigma; E4630) and Rifampicin (Millipore Sigma; R3501) were used as anti-*Mtb* drug control. Solutions of Ethambutol and Rifampicin in distilled water were prepared and stored at 4 °C.

After the day of aerosol infection, female mice were randomly divided into 4 groups (n = 4-8 per group, as indicated in the Results). Feed and water were given *ad libitum*. Treatment with drugs started at 2 weeks (*B6.Sst1S* mice and WT B6 Old), or 4 weeks (for WT B6 mice) post-infection. The drug treatment continued for another 6 weeks (for *B6.Sst1S* mice), 3 weeks (for WT B6 Old) or 4 weeks (for WT B6). Treated mice were administered Ethambutol (100 mg/kg of body weight) by oral gavage, once daily, 5 times a week. Vehicle or Senolytics (Dasatinib: 5 mg/kg, Fisetin: 100 mg/kg, and Quercetin: 50 mg/kg of body weight of mice) were administered by oral gavage, once daily, 3 times a week. The fourth group received both the Ethambutol and Senolytics combination. All drugs were administered in a total volume of 0.2 ml per treatment. Three infected mice were sacrificed at the start of treatment as pretreatment controls. At indicated time points, mice were sacrificed, and the lungs and spleen were harvested for the measurement of bacterial burden, IHC, FACS and histopathology.

### Time to death assay

To examine if the Senolytics administration can improve the survival and body weight, the survival of both male and female *B6.Sst1S* mice (8-10 weeks old) following *Mtb* infection were assessed. Mice were infected via the aerosol route using a Glas-Col aerosolization instrument to deliver a high dose of *Mtb* to the *B6.Sst1S* lungs (approximately 100 CFU). After 3 weeks post-infection, infected mice were randomly divided into 4 groups and the drug was administered as mentioned above, with slight modifications. The drug treatment continued for another 4 weeks, where mice were administered Ethambutol (50 mg/kg of body weight) by oral gavage, once daily, 5 times a week. In the second week, Ethambutol (50 mg/kg of body weight) was combined with Rifampicin (10 mg/kg of body weight) for 1 week. Vehicle or Senolytics (Dasatinib: 5 mg/kg, Fisetin: 100 mg/kg, and Quercetin: 50 mg/kg of body weight of mice) were administered by oral gavage, once daily, 3 times a week. The fourth group received both the Ethambutol and Senolytics combination. All drugs were administered in a total volume of 0.2 ml per treatment. All mice were monitored daily for signs of disease progression, including weight loss, lethargy, hunched posture, and reduced activity. Survival data were analyzed using Kaplan-Meier survival curves to determine the median time to death and assess differences between treatment groups.

### Statistical analysis

Statistical analyses were performed using GraphPad Prism version 10.4.0 software. Normality distribution was first assessed using the Shapiro-Wilk test. For comparisons of lung CFUs, data was log_10_-transformed and checked for normality before the analysis. Statistical significance was determined using unpaired two-tailed Student’s t-tests or Mann Whitney test for two-group comparisons, and one-way or two-way ANOVA or Kruskal-Wallis test (non-parametric test) with post hoc tests for multiple-group comparisons. Kaplan-Meier survival analysis was conducted for time-to-death (TTD) studies, with group differences assessed by the Log-rank (Mantel-Cox) test. In animal experiments, each data point represented a single mouse, and experiments were randomized, though blinding was not implemented. Results are presented as mean with error bars indicating SEM. *p-*values less than 0.05 were considered statistically significant.

### Contact for reagent and resource sharing

All data associated with this study are present in the paper-either in the Results section or in the Supplementary files. Requests for further details on resources, protocol, and reagents can be directed to the corresponding author-Prof. William R Bishai. Raw data generated during this study is provided as a separate Excel file labeled “Source Data file”. The RNA sequencing and Spatial transcriptomics datasets are available on NCBI’s Gene Expression Omnibus (GEO). The *B6.Sst1S* mice used in this study are available for academic and non-commercial research upon request to Prof. Igor Kramnik, under a Material Transfer Agreement.

## Acknowledgements

We gratefully acknowledge the support of the NIH grants AI152688, AI155602, CA006973, and R01HL126066 (to IK). We extend our gratitude to the SKCCC Core centers for their assistance with Histology and Flow cytometry. The funders have no role in study-design, data collection and interpretation, and the decision to submit the work for publication or preparation of the manuscript. We sincerely thank all the members of Prof. Bishai lab and Prof. Kramnik lab, for their valuable suggestions and discussions throughout this project. The schematic figures in this study were created using BioRender (with Academic Lab License).

## Author contributions

Funding Acquisition was contributed by-WB, IK. Conceptualization-WB, IK, SS. Methodology-WB, IK, SS, YMM, BK, SY, LK. Investigation and Formal Analysis-WB, IK, SS. Data curation and Validation-SS, SY, LK. Writing original draft-WB, IK, SS, SY. Review and editing manuscript-WB, IK, SS, YMM, BK, SY, LK.

## Conflict of interest statement

The authors declare that they have no competing or conflicting interests.

## Notes

### Competing Interest Statement

The authors have declared no competing interest.

### Summary of Updates

New data were added. Certain sections were rewritten for better understanding.

