## Supplementary data and Figures for "Elimination of senescent cells with senolytic host-directed therapy reduces tuberculosis progression in mice"

List of authors.

Somnath Shee[1], Yazmin B. Martinez-Martinez[1], Benjamin Koleske[1], Shivraj Yabaji[2], Lester Kobzik[3], Igor Kramnik[2], William Bishai[1,#]

[1] Center for Tuberculosis Research, Johns Hopkins University, School of Medicine, Baltimore, MD, USA.

[2] The National Emerging Infectious Diseases Laboratories (NEIDL), Boston University, Boston, MA, USA.

[3] Cellecta, Inc, Mountain View, CA, USA

Short title: Senolytic therapy for TB

Keywords.

Senescence, Aging, Tuberculosis, Mycobacterium, Senolytics, Dasatinib, Quercetin, Fisetin, Granuloma, Necrosis, SASP, DNA damage

**A**

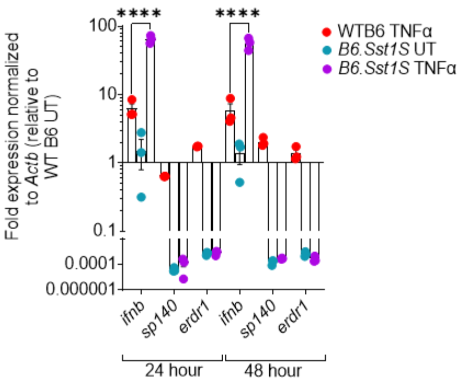

**B**

| | ***B6.Sst1S*(untreated)/ WTB6(untreated) BMDMs** | ***B6.Sst1S*(TNFα)/ WTB6(TNFα) BMDMs#** |
|---|---|---|
| 1.5-fold upregulated | 172 | 778 |
| 1.5-fold downregulated | 89 | 477 |
| 1.5-fold Total DEGs | 261 | 1255 |
| 2-fold upregulated | 21 | 178 |
| 2-fold downregulated | 12 | 50 |
| 2-fold Total DEGs | 33 | 228 |

#: 10 ng/ml TNFα for 18 hours treatment.

**C**

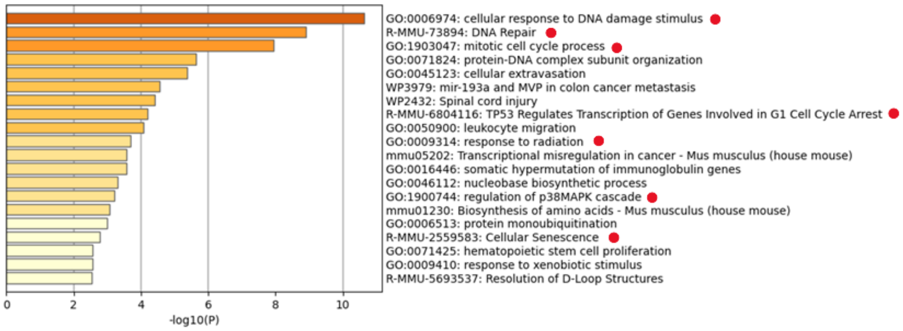

| Network | Annotation |
|---|---|
| MyList | GO:0006974\|cellular response to DNA damage stimulus\|-14.2;GO:0006259\|DNA metabolic process\|-13.1;R-MMU-1640170\|Cell Cycle\|-13.0 |
| MyList_MCODE_ALL | R-MMU-1640170\|Cell Cycle\|-18.8;R-MMU-69278\|Cell Cycle, Mitotic\|-18.4;R-MMU-69481\|G2/M Checkpoints\|-16.9 |
| MyList_SUB1_MCODE_1 | R-MMU-69481\|G2/M Checkpoints\|-20.5;R-MMU-69620\|Cell Cycle Checkpoints\|-17.5;R-MMU-1640170\|Cell Cycle\|-17.2 |
| MyList_SUB1_MCODE_2 | R-MMU-69236\|G1 Phase\|-9.0;R-MMU-69231\|Cyclin D associated events in G1\|-9.0;mmu04218\|Cellular senescence - Mus musculus (house mouse)\|-8.5 |
| MyList_SUB1_MCODE_3 | GO:1901623\|regulation of lymphocyte chemotaxis\|-8.9;GO:0002548\|monocyte chemotaxis\|-8.4;GO:0048247\|lymphocyte chemotaxis\|-8.1 |

**D**

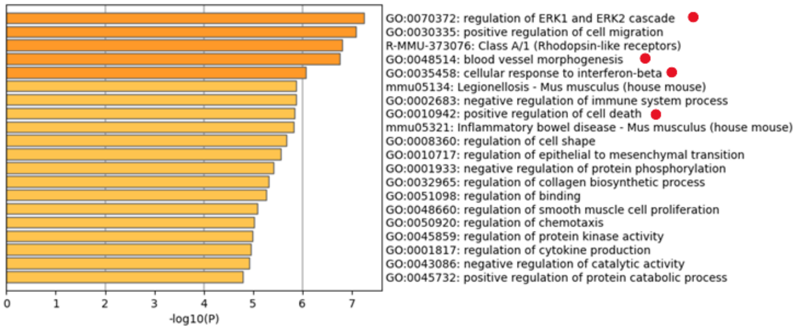

| Network | Annotation |
|---|---|
| MyList | GO:0030335\|positive regulation of cell migration\|-11.5;GO:2000147\|positive regulation of cell motility\|-11.1;GO:0040017\|positive regulation of locomotion\|-10.9 |
| MyList_MCODE_ALL | R-MMU-194315\|Signaling by Rho GTPases\|-7.9;GO:0007264\|small GTPase mediated signal transduction\|-7.9;R-MMU-9716542\|Signaling by Rho GTPases, Miro GTPases and RHOBTB3\|-7.8 |
| MyList_SUB1_MCODE_1 | GO:0007264\|small GTPase mediated signal transduction\|-8.1;R-MMU-9012999\|RHO GTPase cycle\|-6.9;GO:0035970\|peptidyl-threonine dephosphorylation\|-6.9 |
| MyList_SUB1_MCODE_2 | R-MMU-3371453\|Regulation of HSF1-mediated heat shock response\|-9.0;mmu04213\|Longevity regulating pathway - multiple species - Mus musculus (house mouse)\|-9.0;R-MMU-3371556\|Cellular response to heat stress\|-8.5 |
| MyList_SUB1_MCODE_3 | GO:0000278\|mitotic cell cycle\|-7.8;GO:0007052\|mitotic spindle organization\|-6.1;R-MMU-9648025\|EML4 and NUDC in mitotic spindle formation\|-5.9 |

**Figure S1. Transcriptomics evidence suggests emergence of senescence markers in TNFα-treated *B6.Sst1S* BMDMs. (A)** qRT-PCR analysis to show super-induction of *ifnβ* transcripts in TNFα-treated (10 ng/ml) *B6.Sst1S* BMDMs relative to WT B6 BMDMs at the indicated time points. *sp140* and *erdr1x* expression are absent in *B6.Sst1S* mice. *ActB* expression is used as housekeeping control. Error bar represents Mean ±SEM. Data is representative of two independent experiments done in triplicate. Statistical analysis is performed by two-ANOVA with Dunnett's multiple comparisons test. ($p$>0.05: ns, $p$<0.0001: ****).

**(B-D)** Reanalysis of previously published dataset GSE99456. **(B)** Number of differentially expressed genes (DEGs) in indicated comparisons. Briefly, BMDMs from *B6.Sst1S* and WTB6 mice were either left untreated or treated with 10ng/ml TNFα for 18 hours (early time-point) and were subjected to Microarray to understand global transcriptomic changes. (n=3/ group) **(C)** Functional pathway enrichment analysis by Metascape (Express analysis Setting) of DEGs (fold change >1.5) in untreated *B6.Sst1S* BMDMs relative to WT B6 BMDMs revealed enhanced DNA damage response, activation of mitotic checkpoints, and cellular senescence at a basal level, in untreated *B6.Sst1S* BMDMs. **(D)** Functional pathway enrichment analysis by Metascape (Express analysis Setting) of DEGs (fold change >2) in TNFα-treated (10 ng/ml; 18h) *B6.Sst1S* BMDMs relative to analogously treated WT B6 BMDMs revealed ERK1/ERK2 protein kinase cascade, cellular response to IFNβ, epithelial to mesenchymal transition, heat-shock stress, and cell migration pathways to be significantly enriched.

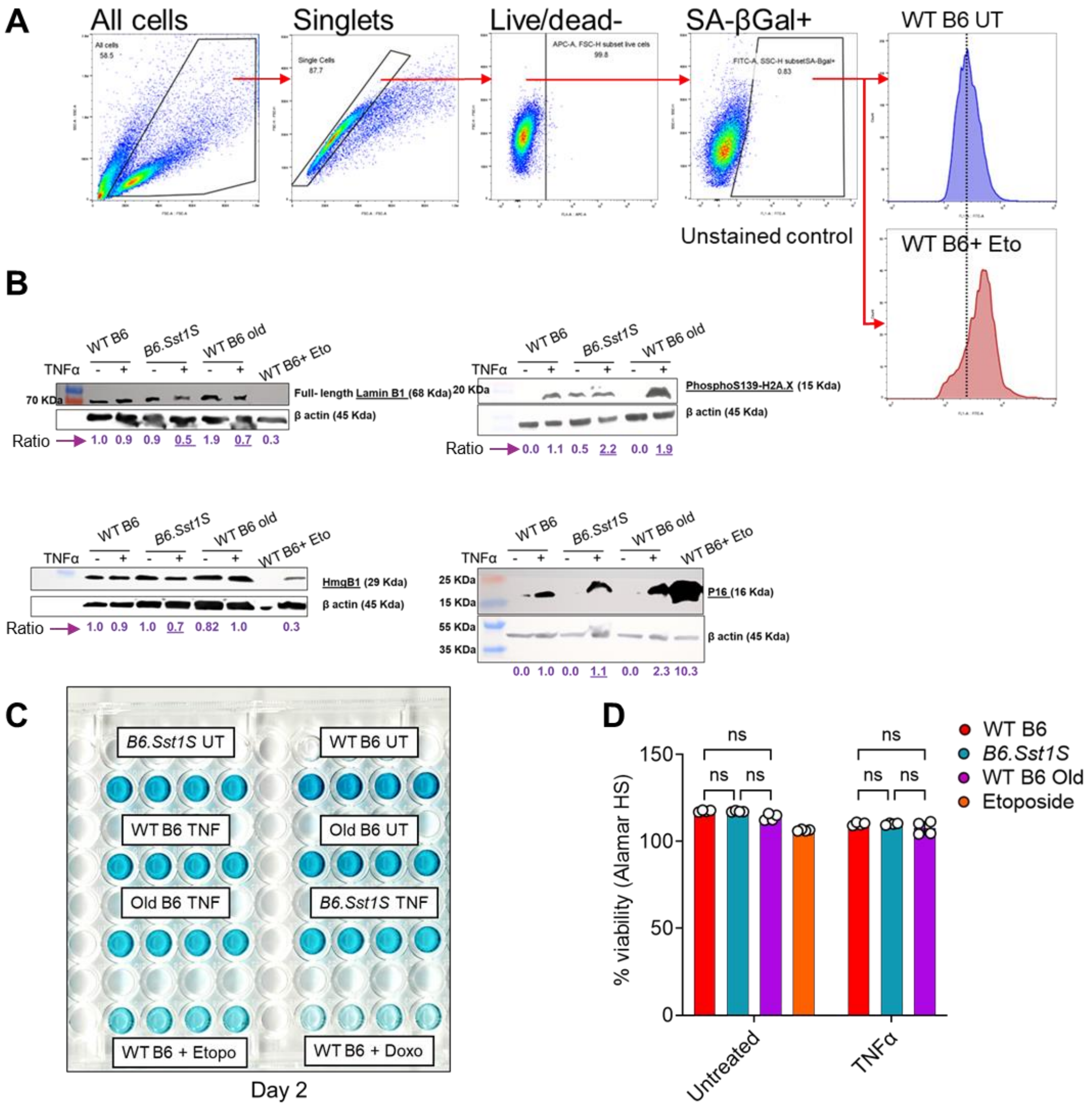

**Figure S2.** Related to Figure 1. **(A)** Representative flow cytometry dot plots and histogram plots to show evaluation of SA-βGalactosidase activity in live BMDMs. WT B6+Eto is positive control where WT B6 BMDMS were treated with etoposide to induce DNA damage-induced senescence. **(B)** Representative Western blots to measure senescence associated protein quantity (Full-length Lamin B1, Hmgb1, γH2A.X and P16) in BMDMs. The number indicates densitometric quantification of bands relative to β-actin loading control. **(C)** Related to Figure 1F**.** Representative images of HRP substrate TMB-color development. The absorbance at 450 nm for the developed color is proportional to the quantity of BrdU incorporated into seeded cells, which is a direct indication of cell proliferation. **(D)** Viability of BMDMs that were either left untreated or treated with TNFα- (10 ng/ml) for 48 h, as measured by a high-sensitivity Alamar blue assay. The fluorescence intensity of each well was measured at excitation/emission of 544/ 590 nm. %viability is relative to the fluorescence of the input well at Day 0. Error bar represents Mean ±SEM. Data is representative of two independent experiments done in triplicate. Statistical analysis is performed by two-ANOVA with Tukey's multiple comparisons test ($p>0.05$: ns, $p<0.0001$: ****).

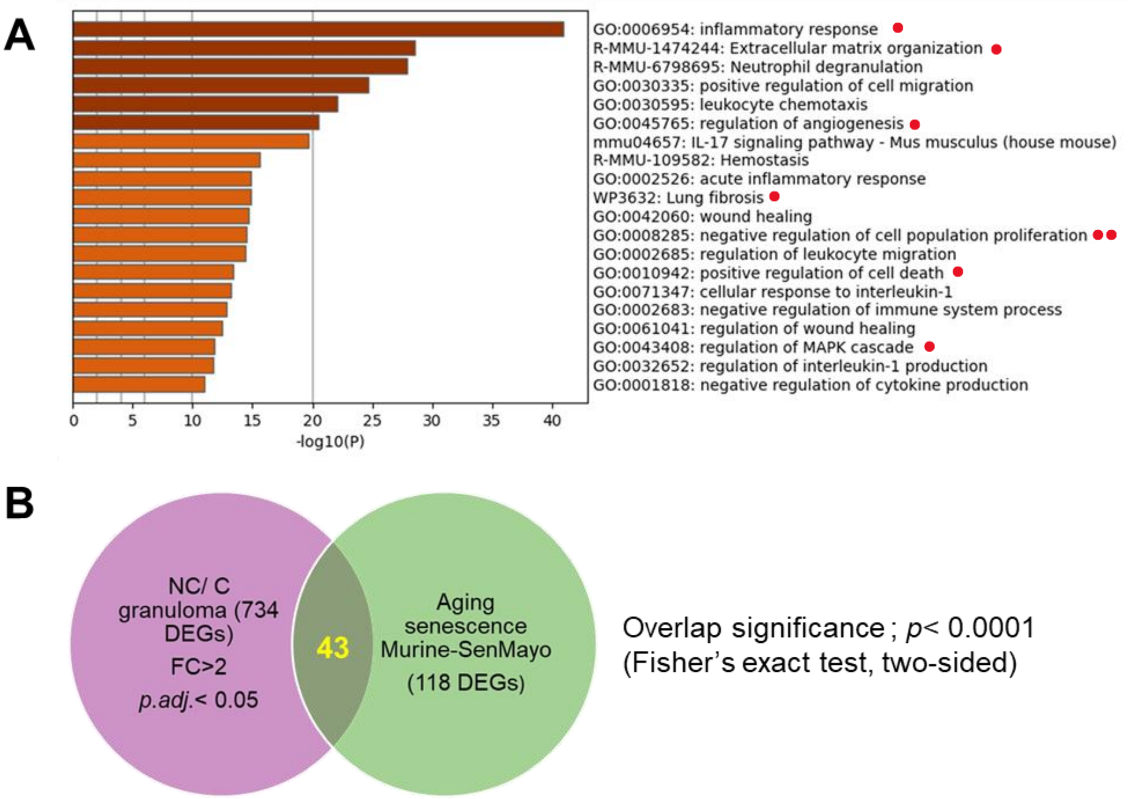

**C**

| | Classification | Gene name |
|---|---|---|
| 1 | Metallo-proteases | Mmp3, Mmp9, mmp12, Mmp13, mmp14, Plat, Plau, Ctsb |
| 2 | Cytokines/ Chemokines | Ccl2, Ccl3, Ccl4, Ccl7, Ccl8, Csf1, Csf2, Cxcl1, Cxcl2, Cxcl3, Cxcl10, Cxcr2, Il1a, Il1b, Spp1, Tnf |
| 3 | Transcriptional Regulators | Ets2, Jun, plaur, Tnfrsf1b, Tnfrsf11b, Cd9 |
| 4 | Intercellular Signal molecule | Angpt1, Angptl4, Esm1, Igfbp4, Igfbp5, Igfbp7 |
| 5 | Negative regulator cell proliferation# | Cdkn1a, Arg1,Arg2 |

#: Not involved in SenMayo gene set.

**Figure S3.** Related to Figure 2A, 2B. **Functional enrichment and senescence-related signatures of upregulated DEGs in cells surrounding NC granulomatous lesions in *B6.Sst1S* mice. (A)** Functional pathway enrichment analysis by Metascape (Express analysis Setting) of upregulated DEGs (419 genes) in NC granulomatous lesions relative to C lesions. Enriched ontology clusters are involved in inflammatory response, extracellular matrix organization, neutrophil degranulation, lung fibrosis, wound healing, and negative regulation of cell population proliferation. GO clusters associated with senescence state are shown with red dots. **(B)** Venn diagram showing overlap with approximately 1/3rd of SenMayo senescence gene set, and **(C)** Table showing the name of these genes and classification based on Saul *et al.*

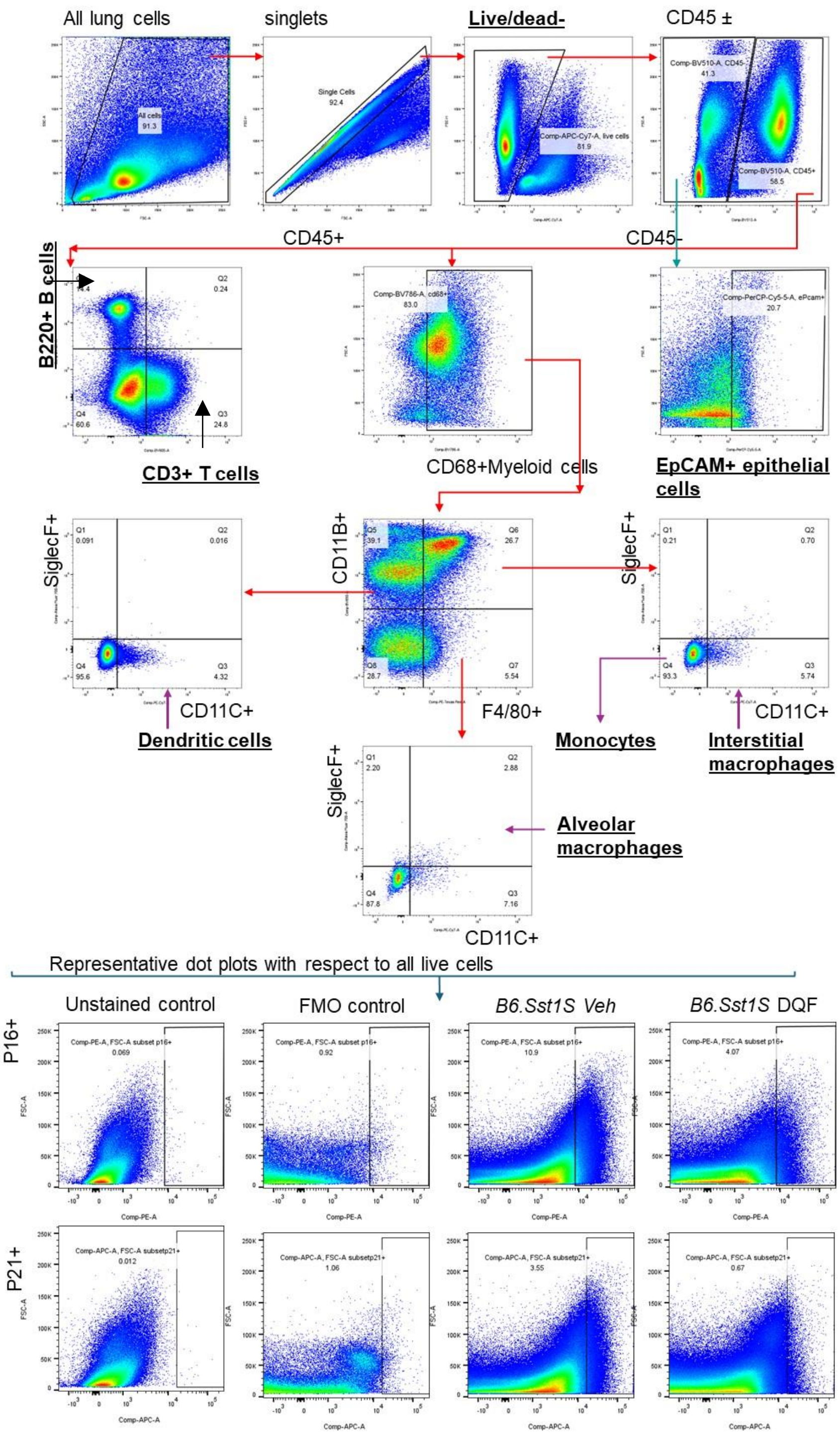

**Figure S4.** Related to Figure 3 and 6. Gating strategy to identify p21+ and p16+ subpopulation in all live cells or different cell populations- myeloid, lymphoid and CD45- cells in the mouse lung. Positively stained cells were gated based on unstained and respective FMO controls using FlowJo_v10.10.0 software.

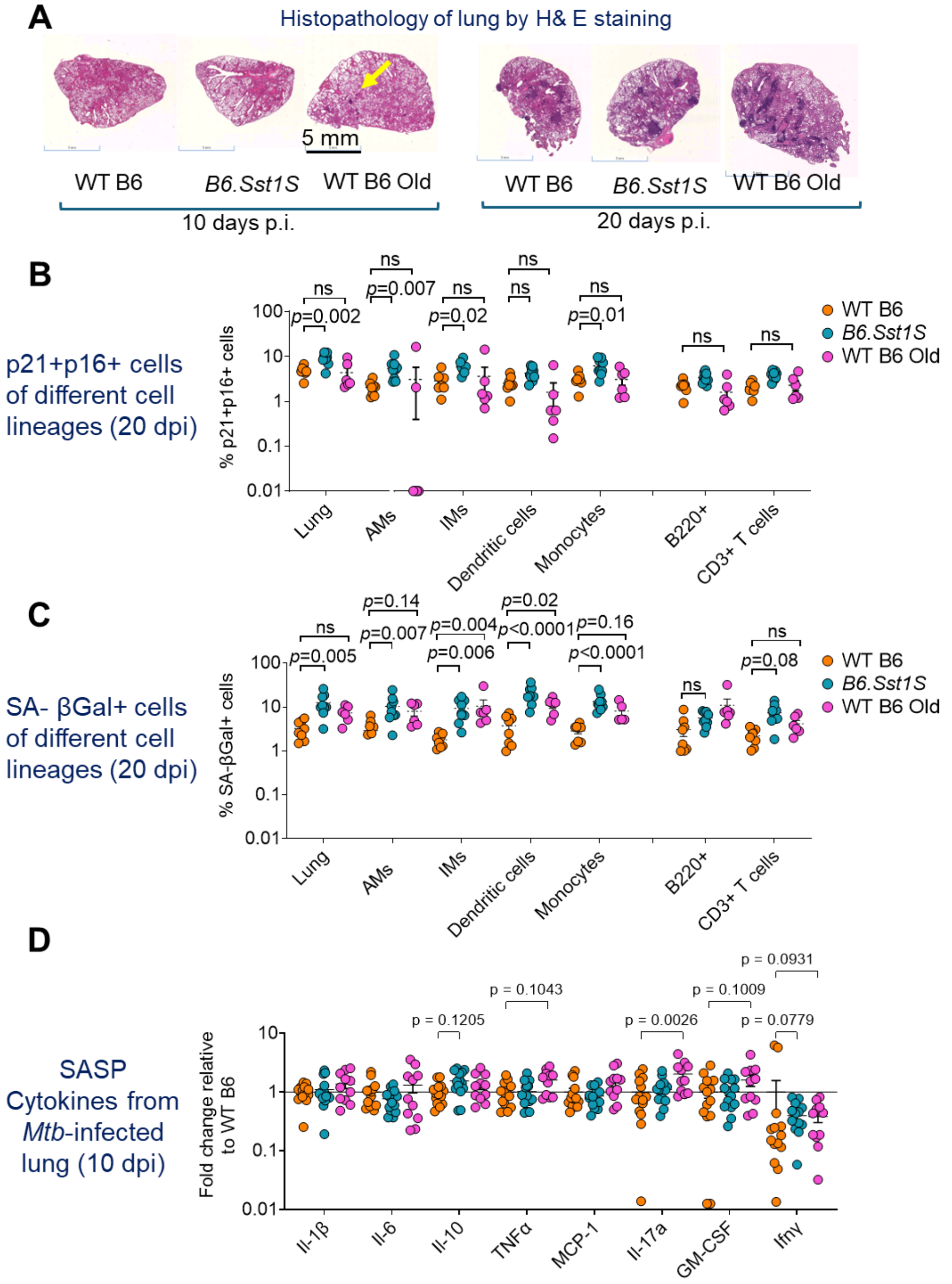

**Figure S5.** Related to Figure 3. **Detection of senescent cell population and SASP cytokines by flow cytometry during early stages of disease progression in *Mtb*-infected mice.** Young WT B6, young *B6.Sst1S*, and old WT B6 mice were aerosol-infected with 55 CFU of *Mtb* H37Rv**. (A)** Representative H&E-stained images of mice lungs at 10 days and 20 days p.i. At 10 days and 20 days p.i., similar histopathology was observed across groups. Early H&E-stained foci (yellow arrow) appeared uniquely in aged WT B6 mice lungs at 10 days p.i. (n=6-9/group). **(B)** %p21+p16+ and **(C)** %SAβGal+ (CellEvent Senescence green+) cells out of all live lung cells and indicated cell types in *Mtb*-infected mice at 20 days p.i., as determined by multicolor flow cytometry. (AMs: Alveolar macrophages, IMs: Interstitial macrophages) The data are means ± SEM. Each data point represents a mouse. Statistics were calculated by two-way ANOVA with Tukey's multiple comparisons test ($p$>0.05: ns). **(D)** Normalized concentration of SASP-cytokines in lung homogenates at 10 dpi and 2 wpi relative to WT B6 mice as measured by a LEGENDplex™ Mouse Inflammation Panel. The data are means ± SEM. Each data point represents a mouse (n=11-14). Statistics were calculated by two-way ANOVA with Dunnett's multiple comparisons test.

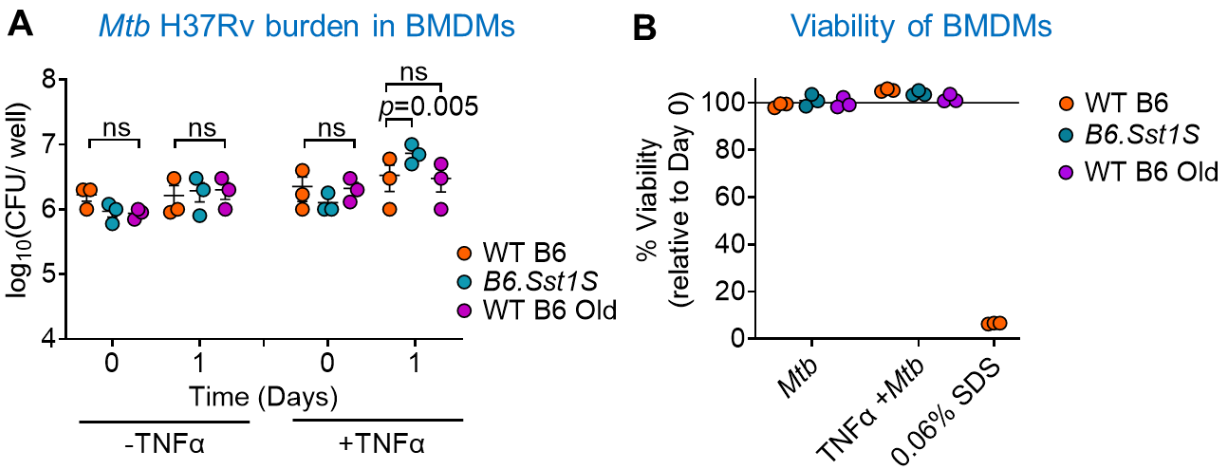

**Figure S6.** Related to Figure 4. **(A)** BMDMs were infected for 4 hours with *Mtb* H37Rv at MOI=1, after overnight incubation with or without TNFα (10 ng/ml). This is followed by removal of extracellular bacteria by extensive washing with warm media. Intramacrophage survival was determined by enumerating CFU at 1-day post-infection. The data are means ± SEM (n=3). Statistics is calculated by one-way ANOVA with Tukey's multiple comparisons test ($p>0.05$: ns). **(B)** In parallel to the sample preparation for RNA-Sequencing experiment, viability of the BMDMs was measured by a high-sensitivity Alamar blue assay. The fluorescence intensity of each well was measured at excitation/emission of 544/ 590 nm. Wells containing only media, and only cells were used as negative and positive control, respectively. %viability is relative to the fluorescence in the positive control well. WT B6 BMDMs treated with 0.06% SDS is used as aa additional negative control.

**A**

| | Drug | MIC 90/Day6 (REMA) | | MBC by growth (Day6 (REMA)+ Day8-10 on 7H11) |
|---|---|---|---|---|
| | | Fluorescence (544/590) | Visual | |
| 1 | Rifampicin | 0.125-0.5 μM | 0.25 μM | 0.25 μM |
| 2 | Ethambutol[#] | 10 μM | 10 μM | 5 μM |
| 3 | Dasatinib | 250-500 μM | 500 μM | **500 μM** |
| 4 | Quercetin | >500-1000 μM | Auto-fluorescence | **>500 μM** |
| 5 | Fisetin | >1000 μM | Auto-fluorescence | **>1000 μM** |
| 6 | DMSO vehicle | Not detected | Not detected | Not detected |

MIC= 90%- Minimum inhibitory concentration at day 6.
#: 88% inhibition.

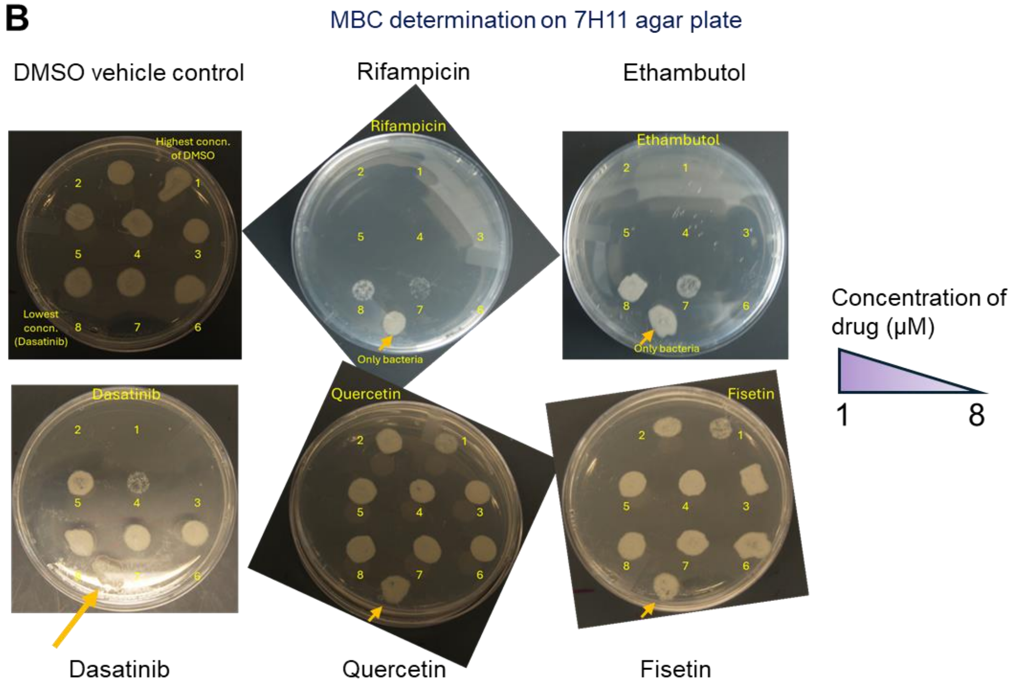

**Figure S7. Senolytics Dasatinib (D), Quercetin (Q) and Fisetin (F) exhibit minimal growth inhibitory effects against *Mtb* H37Rv *in vitro*. (A)** Resazurin Microtiter assay (REMA). Color change from non-fluorescent blue resazurin to fluorescent pink resorufin by cellular metabolic activity indicates reduction of resazurin in the assay. % inhibition of color change with respect to cells-only control is shown in the table. 90% of the inhibition is considered as Minimum inhibitory concentration (MIC). **(B)** 7H11 Agar plate assay for Minimum bactericidal concentration (MBC). 20 μL cells from MIC plate were regrown on 7H11 agar and appearance of bacterial colonies were monitored over time. 1 to 8 corresponds to the highest and lowest concentration of each drug. The golden arrow indicates “only bacteria control without drug”. Q/F had no MBC when measured by growth on drug-containing agar plates. Rifampicin and Ethambutol are used as positive controls. Data shown are representative of two independent experiments done in triplicate.

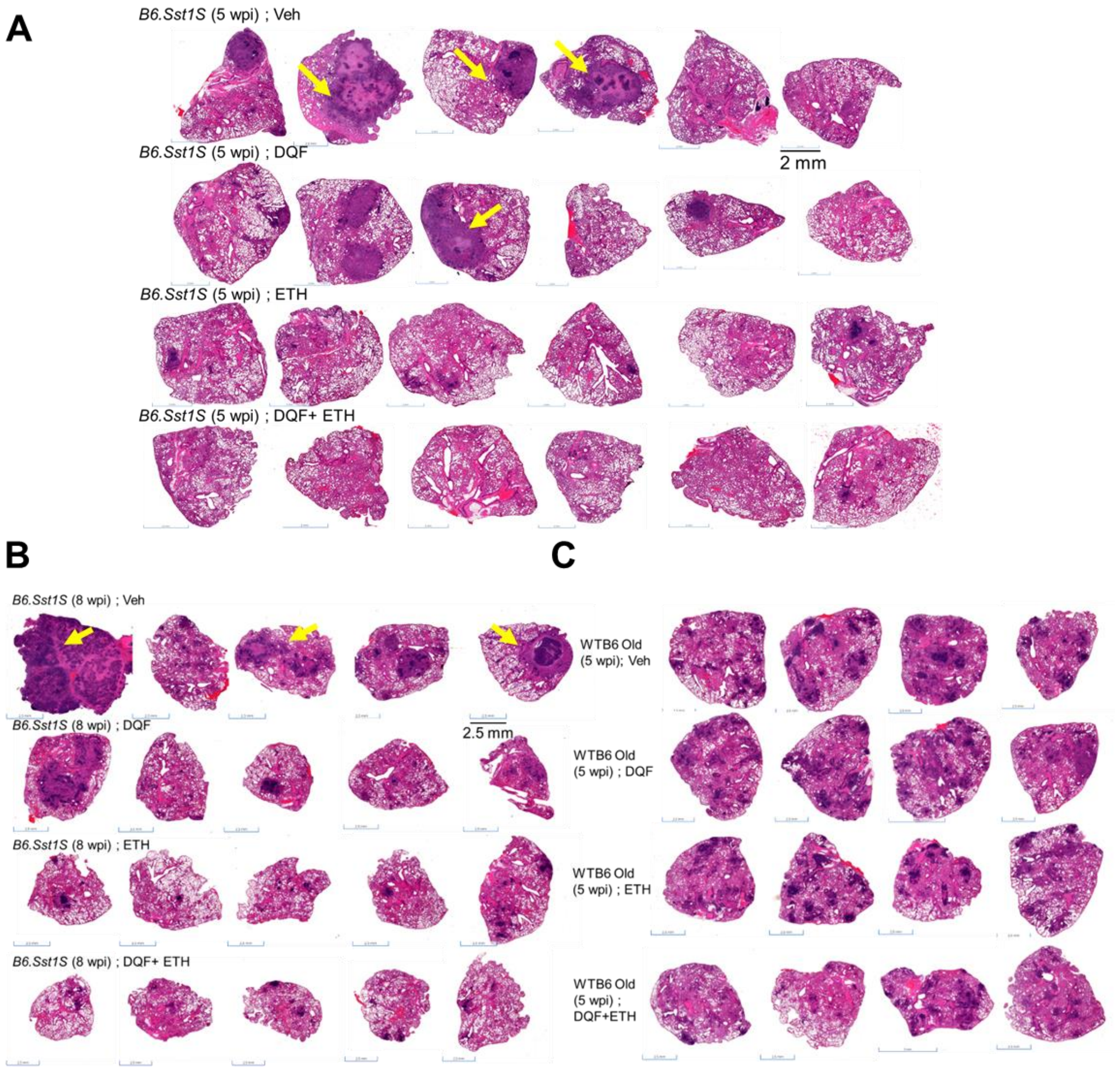

**Figure S8.** Related to Figure 6A. **Senolytics administration alone, and in combination with ethambutol reduced lung pathology in *Mtb* H37Rv infected *B6.Sst1S* and WT B6 old mice.** Histopathology of lungs isolated from 8-12 weeks old female *B6.Sst1S* mice infected with *Mtb* H37Rv at **(A)** 5 weeks p.i. (3 weeks after treatment started; n=6) and **(B)** 8 weeks p.i. (6 weeks after treatment started; n=5). The lungs were formalin fixed, sectioned to 4µm, and stained with hematoxylin and eosin (H&E). Yellow arrows indicate necrotic granulomatous lesions uniquely in *B6.Sst1S* mice. **(C)** H&E-stained lung sections of *Mtb* H37Rv- infected aged WT B6 mice (>72 weeks old) mice at 5 weeks p.i. (3 weeks after treatment started; n=4).

The lung inflammation was lowest in the mice group treated with the combination of DQF+ETH for both *B6.Sst1S* and WT B6 old mice.

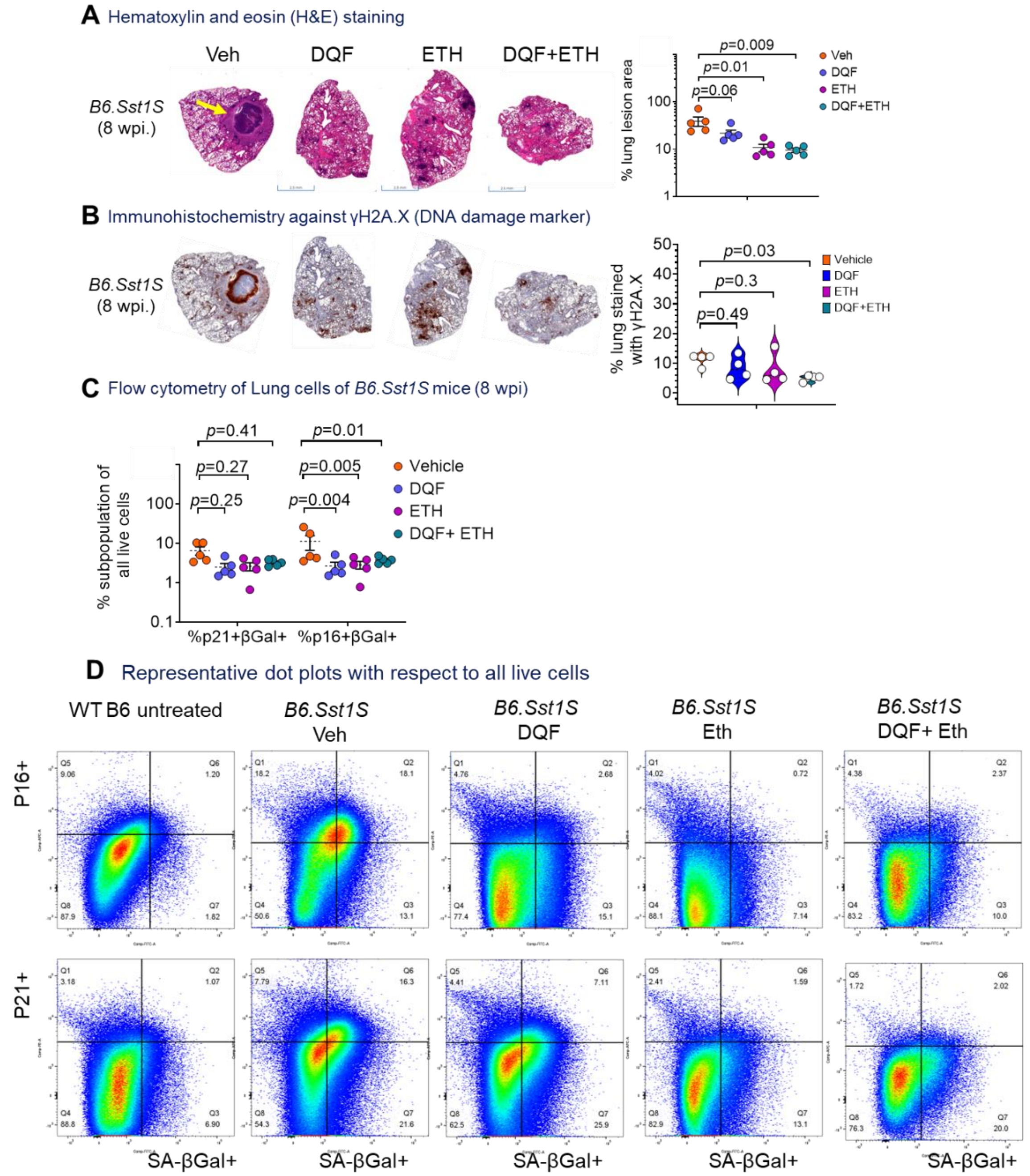

**Figure S9.** Related to Figure 6, 7. **Senolytics administration alone, and in combination with Ethambutol reduced lung pathology and senescence- associated markers in *Mtb* H37Rv infected *B6.Sst1S* mice at 8 wpi. (A)** Representative H&E-stained images of *Mtb*- infected *B6.Sst1S* mice lungs treated with Vehicle or drugs, and respective ImageJ- based quantification. Yellow arrow indicates necrotic granuloma in Vehicle treated *B6.Sst1S* mice at 8 wpi. The data are means ± SEM. Each data point represents a mouse (n=4-5). **(B)** Representative γH2A.X- Immunohistochemistry images of mice lungs after vehicle/ drugs treatment, and respective Violin plots to show ImageJ quantification of γH2A.X-stained area. Each data point represents a mouse. n=4-5 mice/group. Statistical analysis between two groups was done by unpaired two- tailed Student's t test (Normal distribution) or Mann Whitney test. **(C)** %p21+βGal+ and %p16+βGal+ subpopulation observed in all live lung cells of *Mtb*- infected *B6.Sst1S* mice at 8 wpi. (n= 5-8). The data are means ± SEM. Each data point represents a mouse. Statistical analysis was calculated by two-way ANOVA with Dunnett's multiple comparisons test. **(D)** Representative flow cytometry dot plots to show %p21+βGal+ and %p16+βGal+ subpopulation in all live single cells in *Mtb*-infected *B6.Sst1S* mice and WT B6 old mice. Gating for $p16^+$, $p21^+$, and SA-$\beta Gal^+$ events were established using Fluorescence minus one (FMO) controls and reference cells from *Mtb*-infected WT B6 mice (set at 1%).
